# Organ context dominates cell identity across a multi-organ single-cell atlas of grapevine

**DOI:** 10.64898/2026.04.26.720720

**Authors:** Maria-Sole Bonarota, Rosa Figueroa-Balderas, Noé Cochetel, Dario Cantu

**Author notes:** Corresponding author: Dario Cantu. Department of Viticulture and Enology, University of California Davis, Davis, United States.

## Abstract

How transcriptional programs defining cell identity are maintained or remodeled across organ contexts remains a fundamental question in plant biology, particularly in long-lived perennials. We generated a multi-organ single-nucleus transcriptomic atlas of the dwarf grapevine cultivar Pixie, comprising over 220,000 nuclei from nine organs across vegetative and reproductive development, each sampled in two biological replicates. We annotated 46 distinct cell types and anatomical structures, reconstructed developmental trajectories within selected cell types, and inferred gene regulatory networks at cell-type resolution. Organ identity accounted for 65% of regulon activity variance across the atlas compared to only 17% for cell type, demonstrating that organ context is the dominant driver of transcriptional variation in most shared cell types. Broadly distributed cell types including epidermis and xylem parenchyma underwent extensive organ-specific reprogramming, while companion cells maintained organ-independent regulatory programs across all nine organs, representing the stable end of a continuum of transcriptional plasticity. This finding suggests that companion cell identity is buffered against organ context, consistent with their invariant role in phloem loading. This atlas establishes a reference framework for dissecting gene regulatory network organization across organs in a perennial crop and provides a resource for cell-type-resolved functional genomics in grapevine and other woody perennials.

## Introduction

Plant organs are built from a common set of cell types, yet these cells adopt distinct functional states depending on their organ context. Epidermal, vascular, and ground tissues are distributed throughout the plant body, while others, such as pollen and ovules, are confined to reproductive organs and perform specialized roles (Braun, 2022; Harrison Day et al., 2024; Javelle et al., 2011; Shan et al., 2019). This modular organization raises a central question: to what extent is cell identity intrinsic to a cell type versus imposed by its organ environment? In plants, where organogenesis continues throughout life, regulatory programs must repeatedly establish and maintain cell identity in changing organ contexts.

Cell identity is defined by gene regulatory networks in which transcription factors bind specific DNA motifs to drive spatial and temporal expression programs (Badia-i-Mompel et al., 2023; Blanc-Mathieu et al., 2024; Levine and Davidson, 2005). In plants, several transcription factor families establish distinct cell identities, including NAC proteins such as VND6 and VND7 in xylem differentiation, MYB46 and MYB83 in secondary cell wall biosynthesis, R2R3-MYB proteins such as WEREWOLF and GLABRA1 in epidermal patterning, and MADS-box factors in floral organ specification (Honma and Goto, 2001; Kubo et al., 2005; Theißen et al., 2016; Yamaguchi et al., 2008). However, whether these regulatory programs are conserved across organs or reconfigured by organ context remains unresolved at the whole-plant scale, particularly in perennial species with complex tissue organization and secondary growth.

Single-cell and single-nucleus transcriptomics enable gene expression profiling at cellular resolution, allowing the identification of distinct cell populations and transcriptional states within complex tissues (Bonarota et al., 2025; Cantó-Pastor et al., 2024; Illouz-Eliaz et al., 2025; Tenorio Berrío et al., 2025). Single-nucleus approaches are particularly well suited for woody or recalcitrant tissues where protoplasting can bias cell recovery (Grones et al., 2024). When applied across organs, these methods provide a framework to test whether gene regulatory networks defining cell identity are shared or context-dependent (Bonarota et al., 2025; Guo et al., 2025; Kim et al., 2021; Lee et al., 2025). Combined with motif enrichment and regulatory network inference, they also enable identification of candidate regulators of cell identity and tissue specialization (Aibar et al., 2017; Cao et al., 2024; Dorrity et al., 2018).

A key limitation of most existing studies is their focus on single organs. Without a cross-organ framework, it is difficult to distinguish transcriptional programs intrinsic to a cell type from those imposed by organ context. This distinction is central: stable identity across organs implies regulatory buffering, whereas divergence reflects context-dependent reprogramming. Despite this, most plant single-cell studies have been restricted to individual organs in annual models such as Arabidopsis, rice, maize, and tomato (Baumgart et al., 2025; Cantó-Pastor et al., 2024; Denyer et al., 2019; Marand et al., 2021; Shulse et al., 2019; Zhang et al., 2021). Multi-organ atlases are beginning to emerge in these systems (Guo et al., 2025; Lee et al., 2025; Wang et al., 2021; Zhang et al., 2021, 2025), but comparable resources for woody perennials remain scarce. Unlike annual models, perennial species develop complex vascular systems with specialized cell types such as ray parenchyma and vessel-associated cells, which play key roles in carbon allocation, water transport, and whole-plant integration (Chen et al., 2021, 2024; Conde et al., 2022; Li et al., 2021).

Grapevine (*Vitis vinifera* L.) is a perennial crop with high-quality reference genomes and pan-genome assemblies (Cantu et al., 2024; Cochetel et al., 2023; Liu et al., 2024). Its woody growth habit and secondary development generate a structurally complex plant body with diverse vegetative and reproductive organs, encompassing a broad spectrum of cell types, including those specific to perennial systems. These features make grapevine particularly well suited to test how cell identity is maintained or reprogrammed across organ contexts in a long-lived plant. Single-nucleus transcriptomic studies in grapevine have begun to resolve cell type-specific responses within individual tissues (Bonarota et al., 2025a), but a cross-organ framework is still lacking. To overcome the constraint that field-grown vines produce organs at different times, we used Pixie, a dwarf grapevine regenerated from the L1 cell layer of Pinot Meunier (Boss and Thomas, 2002; Cousins, 2012). Pixie carries a dominant mutation in the DELLA gene *VvGAI1* that partially disrupts gibberellin signaling, resulting in dwarfism and continuous conversion of tendrils into inflorescences, enabling year-round flowering and fruiting under greenhouse conditions. Although Pixie differs in architecture and fruit size from standard cultivars, the major cell types profiled here are conserved, supporting the broader relevance of this atlas.

Here, we present a multi-organ single-nucleus transcriptomic atlas of grapevine comprising over 220,000 nuclei from nine organs spanning vegetative and reproductive development. The atlas includes dormant buds and berries at multiple developmental stages, capturing transitions between dormancy, active growth, and fruit development, and encompasses 46 annotated cell types and anatomical structures. We show that cell types shared across organs span a continuum from context-dependent to context-independent transcriptional programs. Notably, companion cells maintain a stable regulatory signature across diverse organs, whereas xylem parenchyma and other shared cell types exhibit pronounced organ-specific divergence. We further identify transcription factors and cis-regulatory motifs associated with specific cell populations, providing candidate regulators of cell-type identity and tissue specialization. Together, this atlas establishes a framework for linking gene regulatory networks to cell identity across organs in a perennial species whose developmental and vascular complexity extends beyond that captured in annual models.

## Material and Methods

### Plant material and experimental design

Nine organs (roots, dormant buds, green stems, young and old leaves, pre-anthesis flowers, and green, harvest and postharvest berries) from the grapevine Pixie FPS clone 01 were sampled in duplicate to construct a single-nucleus gene expression atlas. The samples used were identical to those described in Paineau et al. (2025). All samples were collected at 10:00 a.m. Dormant buds, green stems, young and old leaves and pre-anthesis flowers were harvested from Pixie plants grown in greenhouse with 16h photoperiod, with temperatures ranging from 16C (night) to 23C (day), and humidity at ∼60%. The 5-cm distal region of the root system was harvested from cloning green cuttings in a plant cloner with 10% Hoagland fertilized water in the same environmental conditions. Berry samples included three developmental stages: green berry (hard, green, pre-veraison), harvest berry (colored, soft, ripe), and postharvest berry (very soft, over-ripe). Two biological replicates were used for each organ, and samples from five different plants were pooled into one biological replicate.

### Nuclei isolation, library preparation and sequencing

Nuclei isolation and single-nuclei RNA sequencing (snRNA-seq) libraries for root, bud, stem, leaf and flower organs were generated by SolusCell (Alachua, FL, USA) using methods optimized for plant tissues. Briefly, nuclei were isolated from flash frozen tissue as described previously (Conde et al., 2022), followed by purification using proprietary methods. Nuclei were used to generate snRNA-seq library following the GEXSCOPE Single Nucleus RNA Library Kit V2 protocol from Singleron Biotechnologies (Cologne, Germany). These services were performed on a fee-for-service basis; SolusCell had no role in study design, data interpretation, or manuscript preparation. Single-nuclei isolation from grapevine berries was modified by Bonarota et al., (2025). The protocol was optimized using a Plant Nuclei Isolation/Extraction Kit (CelLytic™ PN, Sigma). Around 1.5 g of frozen deseeded berries were finely ground. The sample was transferred in a 50-mL Falcon tube, the nuclei isolation buffer (NIB) was added to reach 25 mL volume, slowly inverted two times and incubated on ice for 15 min. Slow inversions were repeated every 3 min. The sample was filtered using a 20 μm steriflip vacuum filter and then centrifuged at 5000 rpm (4696 RCF) for 10 min. After decanting the supernatant, NIB-A was added (5 mL) to the pellet to resuspend. Triton 10% (250 μL) was added to the solution and incubated for 5 min. The solution was filtered using 30 μm Miltenyil filters and added to 5 mL, 1.5 M sucrose cushion. After centrifugation at 5000 rpm for 10 min, the supernatant was carefully removed, and the pellet was washed twice with phosphate-buffered saline 1X and 2% bovine serum albumin (3.5 mL), through centrifuging at 5,000 rpm for 5 min. Finally, the pellet was resuspended in 50 μL PBS 1X + 2% BSA and filtered through a PBS-pre-cleaned in-house cotton column to remove debris (Lee and Lin, 2005). Nuclei quality was assessed through DAPI and AO/PI staining at 20x and 40x magnification. Nuclei quantification was performed with a fluorescent cell counter LUNA-FL (Logos Biosystem, Inc., South Korea). Final nuclei concentrations were between 600 and 700 nuclei μL^-1^. Single nuclei were partitioned and barcoded using the Chromium Next GEM Single Cell 3’ Reagent Kit V4.0 (10x Genomics). Sequencing was performed through the UC Davis Genome Center on an Illumina NovaSeq X 25B flow cell, using 150-bp paired-end sequencing, with an average sequencing depth of 40,000 reads per nucleus.

### snRNA-seq analysis

For the alignment of snRNA-seq data, a custom reference genome was generated by concatenation of the genomes of *V. vinifera* cv. Pixie clone FPS01 (https://www.grapegenomics.com/pages/VvPixie/; Paineau et al., 2025), grapevine chloroplast ( Arroyo-Garcia et al., 2006) and mitochondria (Goremykin et al., 2009). Reads were mapped using alevin-fry (He et al., 2022) using the simpleaf v0.19.4 framework (He and Patro, 2023). The average mapping rate was 87.13 ± 7.73 % (listed per sample in **Table S1**). The count matrix was extracted with loadfry function from the fishpond package v2.16.0 (Zhu et al., 2019) in R v4.5.3. Empty barcodes were filtered out using DropletUtils v1.30.0 (Lun et al., 2018). The following analysis was performed with the Seurat package v5.3.0 (Satija et al., 2015) for each organ separately. Low-quality nuclei with fewer than 200 and higher than 3,000 genes and with more than 5% of reads mapping to the mitochondrial and chloroplast genomes were discarded. Genes expressed in less than three nuclei were discarded. To account for batch effect between samples, we integrated the data within organs using FindIntegrationAnchors to identify the anchors and IntegrateData (with normalization.method = “SCT”). To assess potential batch effects across organs associated with the use of different library preparation platforms, we evaluated transcriptional variation within cell types shared between berry and non-berry tissues. Specifically, we performed principal component analysis (PCA) on gene expression profiles for six cell types detected in both datasets: epidermis, chlorenchyma, companion cells, vascular cells, phloem parenchyma, and xylem parenchyma. For each of the first 20 principal components (PCs), we fitted three separate linear models to quantify the proportion of within-cell-type variance associated with: (1) organ/stage identity, (2) library preparation platform (GEXSCOPE versus 10x Genomics), and (3) organ/stage identity restricted to organs profiled using the GEXSCOPE platform. The coefficient of determination (R²) was calculated for each model and averaged across the 20 PCs to obtain an overall estimate of the variance explained by each factor. This analysis allowed us to compare the contribution of biological organ/stage identity with that of library platform and to assess whether organ/stage-associated variation remained evident independently of the inclusion of berry samples generated using the 10x Genomics platform.

Data were normalized using SCTransform, followed by principal component analysis (PCA) using RunPCA (with 5,000 variable genes and 30 to 100 principal components (PCs) depending on the complexity of the organ), construction of K-nearest neighbors with FindNeighbors (k-param = 20), and calculation of shared nearest neighbors using FindClusters (resolution = 1 or 2, depending on the complexity of the organ). Data were visualized with non-linear dimensionality reduction uniform manifold approximation and projection (UMAP; Healy and McInnes, 2024) using the function RunUMAP. For cluster annotation, a list of marker genes was created using the function FindAllMarkers on raw expression data (with logfc.threshold = 0.1, min.pct = .05, only.pos = T, test.use = ‘wilcox’). Cell type markers were filtered to p_val_adj < 0.05. Cell type marker genes were visualized with the DotPlot function, using parameter scale = T, which scales the average expression values across all genes to have a mean of 0 and variance of 1 within each gene. Whenever the cell type marker list was not sufficient to assign annotation, we recurred to the past grapevine bulk tissue transcriptomic literature (e.g. flower stigma, pollen, and anther, berry exocarp, mesocarp, and bud dormant and proliferating meristem; see **Supplementary Text S1**). To validate cell type markers, genes were ranked along a continuum from broadly expressed to highly cell-type-specific using τ specificity index (Yanai et al., 2005). Additionally, the gene expression of main cell-type markers used in this study was compared in *Arabidopsis thaliana* (Lee et al., 2025). Orthologs were identified using AEGIS (Navarro-Payá et al., 2026), and only high-and medium-confidence orthologs were retained.

### Development of a grapevine single-nucleus transcriptomic atlas web portal

An interactive web portal was implemented using Claude Anthropic tool as a static, client-side single-page application (vanilla JavaScript ES modules, Plotly.js for visualization), requiring no backend server. Per-cell-type average expression, percent of cells expressing each gene, and z-scored activity for both genes and inferred transcription-factor regulons were computed from the Seurat objects for each organ and for the combined all-organ dataset. Summary statistics were then pre-exported to static JSON files via an R export pipeline, together with cell coordinates (UMAP embeddings) and cell-type annotations. Gene identifiers were cross-referenced between the Pixie genome assembly and the PN40024 v5.1 (T2T) reference genome using MCScanX and AEGIS suite (Navarro-Payá et al., 2026; Tang et al., 2024), allowing users to query and display genes under either nomenclature. The portal is deployed as a static site and embedded via iframe on www.grapegenomics.com (https://www.grapegenomics.com/pages/atlas.php).

### Analysis of gene regulatory networks in cell types and organs

To understand and predict cell type-specific gene regulatory networks (GRNs) across cell types and organs, we used Motif-Informed Network Inference based on single-cell Expression data (MINI-EX) V2 (Staut et al., 2026) with default parameters. MINI-EX2 is designed to integrate single-cell RNA sequencing data (scRNA-seq) with TF binding motif information, inferring high-quality GRNs for each cell type, and prioritize most relevant regulators using network statistics and gene ontology.

The transcription factor (TF) list and corresponding family was downloaded from PlantTFDB (http://planttfdb.gao-lab.org/index.php) (Jin et al., 2016). The TF motif information was obtained from JASPAR (https://jaspar.genereg.net/downloads) (Castro-Mondragon et al., 2022). TF orthology from PN40024 to Pixie grapevine was carried out using MCScanX (Wang et al., 2024) from the JCVI v.1.0.9 toolkit (Tang et al., 2024) and validated by AEGIS suite (Navarro-Payá et al., 2026). Motif occurrences across the Pixie genome were identified with Find Individual Motif Occurrences (FIMO) (Grant et al., 2011), within the suite meme-5.5.9 (Bailey et al., 2015).

We used the top 750 upregulated genes per cluster and filtered out regulons (which consist of the TFs and their putative targets) of single-cell clusters where the TF is expressed in less than 1% of the cells. Due to low number of nuclei in some cell types in buds and roots, we merged root tip cells (columella and peripheral cells), vascular cells (phloem and xylem parenchyma, companion cells and developing sieve elements) and epidermal cells (e.g., exodermis and hypodermis) in the same clusters. In bud and root organs, we used the top 1000 upregulated genes per cluster and did not filter out lowly expressed regulons of single-cell clusters to account for low number of nuclei.

After identifying the regulons within each organ, the regulons’ activities were computed with AUCell 1.28.0 (Aibar et al., 2017) in the whole atlas and across all organs and cell types. AUCell calculates the enrichment of the regulon as an area under the recovery curve (AUC) across the ranking of all genes in a particular cell, whereby genes are ranked by their expression value. This method is independent of the gene expression units and the normalization procedure, and it can easily be applied to bigger data sets. The average activity scores of all identified regulons per organ, cell type and replicate were analyzed across the whole atlas dataset through the linear mixed model specified by Regulon Activity ∼ Organ * Cell type + (1 | Replicate), to split total variance explained by organ, cell type, and noise (residual variance). Regulon activities were visualized using UMAP, and Wilcoxon test (adjusted p-value < 10^-8^) was used to determine whether a regulon was overexpressed in a specific cell type or organ compared to all other cell types or organs.

### Cell type specific developmental trajectory analysis

To identify developmental trajectories within cell types, clusters were redefined to identify cell types at different developmental stages, such as immature to germinating pollen, green to mature berry exocarp and mesocarp, and young to senescent leaf epidermis and mesophyll. Cell type specific developmental trajectory analysis was performed using Monocle 3 (Trapnell et al., 2014). Briefly, nuclei from each specific cell type were subset in a new Seurat object. Data was normalized, clustered, and UMAP-based plots were built as described above, using a resolution of 0.6. The trajectory graph was learnt by the learn_graph function in Monocle 3. The lowest pseudotime was assigned to the nuclei group with the lowest progress through the biological process (e.g. immature pollen, green berry and young leaf). Differentially expressed genes were identified using the graph_test function in Monocle 3, and only genes with status= “OK” and q-value < 0.05 were considered differentially expressed.

### Pseudobulk snRNA-seq analysis across grapevine organs

Annotated data from each organ were merged into one dataset to form a comprehensive grapevine single-nucleus transcriptomic atlas. Merged data was normalized and the UMAP was built as described above. The raw counts from epidermis, xylem and phloem parenchyma, and companion cell types from each organ were pooled into one dataset each, and normalization and visualization analysis was repeated as explained in the precedent paragraph. Organ-dependent gene expression in epidermis, xylem and phloem parenchyma and companion cells was assessed using a likelihood ratio test (LRT) in DESeq2 (Love et al., 2014), comparing a full model including organ identity to a reduced intercept-only model. Genes with FDR < 0.05 were considered differentially expressed between organs. For significant genes, variance-stabilized expression values were used to compute mean expression per organ.

### Disclosure of AI Use

Claude (Anthropic) was used to develop the interactive single-nucleus atlas explorer (September 2026), as well as to review grammar and style and suggest wording for selected passages during the preparation of the manuscript. All suggestions were critically evaluated by the authors, and the scientific content, interpretation, and conclusions are the sole responsibility of the authors.

## Results

### Single-nucleus transcriptomic atlas of grapevine

Intact nuclei were isolated from old and young leaves, berries at three developmental stages, pre-anthesis flowers, green stems, dormant buds, and distal 5-cm root ends of the grapevine cultivar Pixie, with two biological replicates per organ and developmental stage (18 samples total; **Fig. 1A**). After quality filtering, we retained an average of 12,332 ± 12,483 nuclei per sample (mean ± SD; **Fig. 1B**), reflecting substantial variation in recovery across organs, from 27,357 ± 7,252 nuclei per sample in young leaves to 1,572 ± 1,444 in postharvest berries (**Fig. 1B**; **Table S1**). Transcript capture efficiency also differed among organs, with the highest number of reads and genes per nucleus in green berries (4,574 ± 435 reads; 434 ± 22 genes) and the lowest in roots (506 ± 19 reads; 261 ± 2 genes) (**Fig. 1B**; **Table S1**). Pseudobulk gene expression profiles were highly correlated across biological replicates (Pearson r > 0.9), both globally and within individual cell types, indicating minimal transcriptional variability between replicates (**Fig. S1**).

**Figure 1.**
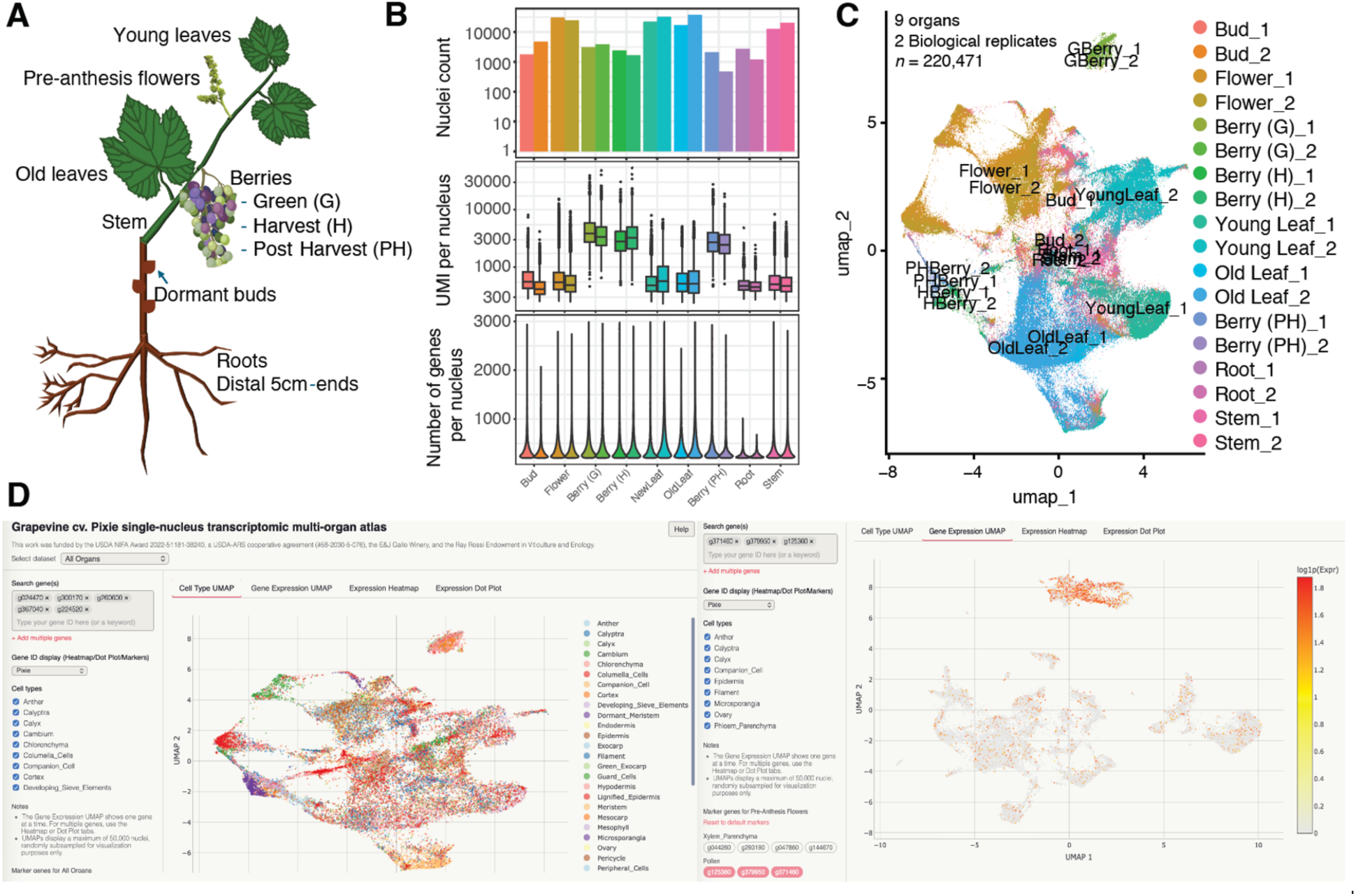
Grapevine multi-organ single-nucleus transcriptomic atlas. (A) Schematic representation of the nine organs profiled in this study, including roots, dormant buds, green stems, young and old leaves, pre-anthesis flowers, and berries at three developmental stages (green, harvest, and postharvest). (B) Quality metrics per sample: total nucleus count (top), UMI per nucleus (middle), and genes detected per nucleus (bottom), shown across all 18 samples. (C) UMAP embedding of 220,471 nuclei from nine organs and two biological replicates, colored by sample identity, showing that nuclei cluster primarily by organ of origin. (D) Web browser of the grapevine single-nucleus atlas showing the integrated organs and an example of gene expression UMAP for pollen marker gene in the flower dataset.

All snRNA-seq datasets were integrated into a single atlas comprising 220,471 nuclei across organs and two biological replicates per organ (**Fig. 1C**). Dimensionality reduction revealed that nuclei clustered primarily by organ, with photosynthetic tissues (leaves and green berries), reproductive tissues (flowers), and structurally specialized organs (roots, buds, and stems) occupying distinct regions of the UMAP (**Fig. 1C**). Biological replicates were intermingled within each organ cluster, confirming successful batch integration. Library preparation technology (GEXSCOPE and 10x Genomics) explained between 0.5 % and 9.8 % variance, whereas organ identity explained between 15.5 % and 51.7 % (**Fig. S2**), showing that platform effects were consistently small relative to organ identity.

The web browser (available at https://www.grapegenomics.com/pages/atlas.php) allows interactive exploration of the Pixie multi-organ single-nucleus atlas across nine organs and developmental stages (dormant bud, pre-anthesis flower, berry, leaf, root, stem) individually or jointly ("All Organs"), as well as the inferred gene regulatory programs (regulons), described below. Users can search for genes by Pixie ID, PN40024 v5.1 (T2T) ID, or functional-description keyword, select one or more genes, and visualize their expression as cell-type UMAPs, expression heatmaps, or dot plots, filterable by cell type. Each organ is pre-loaded with a curated panel of marker genes (**Fig. 1D**), and transcription-factor regulons can be browsed by their target-gene sets. This provides the community with direct, code-free access to the atlas’s expression data without needing to reprocess the underlying sequencing data.

### Cell type identification and annotation

Cell types were defined using Louvain-based clustering performed separately for each organ, with clusters annotated using statistically significant marker genes (log_2_FC > 2; p-adj < 0.05). Across nine organs, we identified 46 distinct cell types and anatomical structures supported by marker gene expression (**Table 1**; **Fig. 2**, **Tables S2-S7**). Detailed marker gene annotations for all cell types and anatomical structures, including gene identifiers and supporting references, are provided in **Tables S2-S7** and **Supplementary Text S1**. Gene markers were validated using τ specificity index (closer to 1 for more specific markers; listed in **Tables S2-S7**). Detailed gene identifiers from the Pixie genome, syntenic gene identifiers in the PN40024 genome, and corresponding gene functional annotation are listed in **Table S8**.

**Figure 2.**
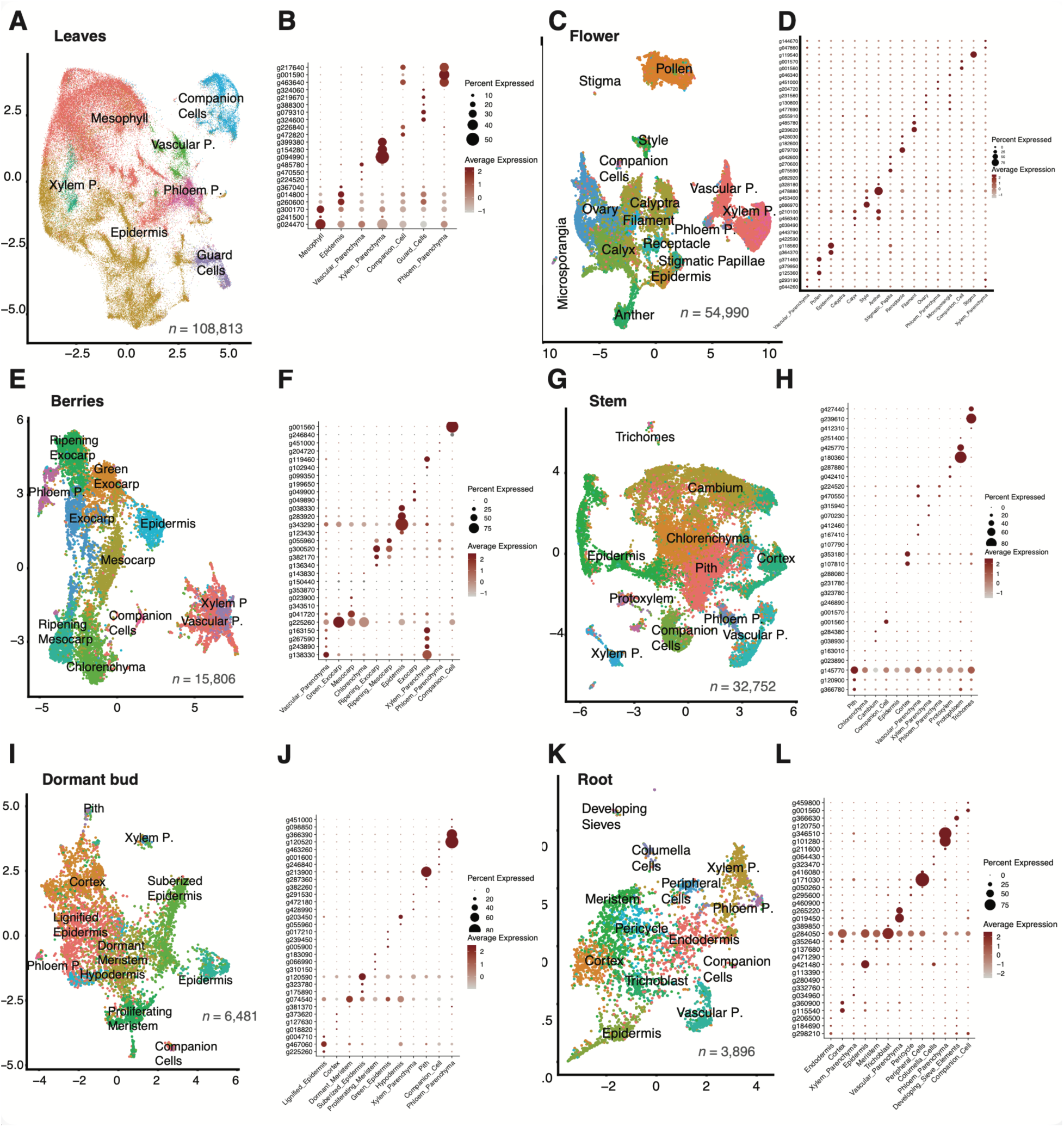
Cell type annotation of a multi-organ grapevine single-nucleus transcriptomic atlas. UMAP-based clustering and marker gene expression used for cell type annotation across nine organs of grapevine cultivar Pixie. For each organ, UMAP embeddings of single-nucleus transcriptomes (left panels) are shown alongside dot plots of pseudobulk expression of marker genes across annotated cell types (right panels). Organs include leaves at two developmental stages (A-B), pre-anthesis flowers (C-D), berries at three developmental stages (E-F), stems (G-H), dormant buds (I-J), and roots (K-L). Dot plots display percent expressed and average expression of main selected marker genes used to define each cell type. Two biological replicates were included for each organ and developmental stage.

**Table 1.** Cell types and anatomical structures in each organ, and relative number of nuclei per cell type per organ.

|  | **Flower** | **Stem** | **Berry (G)** | **Berry (H)** | **Berry (PH)** | **Root** | **Bud** | **NewLeaf** | **OldLeaf** |
|---|---|---|---|---|---|---|---|---|---|
| **Anther** | 2365 | 0 | 0 | 0 | 0 | 0 | 0 | 0 | 0 |
| **Calyptra** | 9431 | 0 | 0 | 0 | 0 | 0 | 0 | 0 | 0 |
| **Calyx** | 7625 | 0 | 0 | 0 | 0 | 0 | 0 | 0 | 0 |
| **Cambium** | 0 | 6980 | 0 | 0 | 0 | 0 | 0 | 0 | 0 |
| **Chlorenchyma** | 0 | 4955 | 2146 | 90 | 76 | 0 | 0 | 0 | 0 |
| **Columella_Cells** | 0 | 0 | 0 | 0 | 0 | 90 | 0 | 0 | 0 |
| **Companion_Cell** | 254 | 2027 | 60 | 31 | 6 | 24 | 72 | 3365 | 4939 |
| **Cortex** | 0 | 3650 | 0 | 0 | 0 | 461 | 1147 | 0 | 0 |
| **Developing_Sieves** | 0 | 0 | 0 | 0 | 0 | 34 | 0 | 0 | 0 |
| **Dormant_Meristem** | 0 | 0 | 0 | 0 | 0 | 0 | 1122 | 0 | 0 |
| **Endodermis** | 0 | 0 | 0 | 0 | 0 | 520 | 0 | 0 | 0 |
| **Epidermis** | 1625 | 4391 | 532 | 256 | 136 | 437 | 404 | 20882 | 19872 |
| **Exocarp** | 0 | 0 | 219 | 623 | 279 | 0 | 0 | 0 | 0 |
| **Filament** | 982 | 0 | 0 | 0 | 0 | 0 | 0 | 0 | 0 |
| **Green_Exocarp** | 0 | 0 | 1650 | 107 | 44 | 0 | 0 | 0 | 0 |
| **Guard_Cells** | 0 | 0 | 0 | 0 | 0 | 0 | 0 | 2604 | 2378 |
| **Hypodermis** | 0 | 0 | 0 | 0 | 0 | 0 | 314 | 0 | 0 |
| **Lignified_Epidermis** | 0 | 0 | 0 | 0 | 0 | 0 | 1357 | 0 | 0 |
| **Meristem** | 0 | 0 | 0 | 0 | 0 | 784 | 0 | 0 | 0 |
| **Mesocarp** | 0 | 0 | 1037 | 315 | 245 | 0 | 0 | 0 | 0 |
| **Mesophyll** | 0 | 0 | 0 | 0 | 0 | 0 | 0 | 21322 | 21881 |
| **Microsporangia** | 398 | 0 | 0 | 0 | 0 | 0 | 0 | 0 | 0 |
| **Ovary** | 8112 | 0 | 0 | 0 | 0 | 0 | 0 | 0 | 0 |
| **Pericycle** | 0 | 0 | 0 | 0 | 0 | 212 | 0 | 0 | 0 |
| **Peripheral_Cells** | 0 | 0 | 0 | 0 | 0 | 108 | 0 | 0 | 0 |
| **Phloem_Parenchyma** | 764 | 804 | 42 | 189 | 210 | 66 | 69 | 2407 | 1788 |
| **Pith** | 0 | 5468 | 0 | 0 | 0 | 0 | 78 | 0 | 0 |
| **Pollen** | 8380 | 0 | 0 | 0 | 0 | 0 | 0 | 0 | 0 |
| **Proliferating_Meristem** | 0 | 0 | 0 | 0 | 0 | 0 | 688 | 0 | 0 |
| **Protophloem** | 0 | 282 | 0 | 0 | 0 | 0 | 0 | 0 | 0 |
| **Protoxylem** | 0 | 502 | 0 | 0 | 0 | 0 | 0 | 0 | 0 |
| **Receptacle** | 1021 | 0 | 0 | 0 | 0 | 0 | 0 | 0 | 0 |
| **Ripening_Exocarp** | 0 | 0 | 39 | 846 | 495 | 0 | 0 | 0 | 0 |
| **Ripening_Mesocarp** | 0 | 0 | 6 | 706 | 453 | 0 | 0 | 0 | 0 |
| **Stigma** | 80 | 0 | 0 | 0 | 0 | 0 | 0 | 0 | 0 |
| **Stigmatic_Papilla** | 1037 | 0 | 0 | 0 | 0 | 0 | 0 | 0 | 0 |
| **Style** | 1231 | 0 | 0 | 0 | 0 | 0 | 0 | 0 | 0 |
| **Suberized_Epidermis** | 0 | 0 | 0 | 0 | 0 | 0 | 1129 | 0 | 0 |
| **Trichoblast** | 0 | 0 | 0 | 0 | 0 | 257 | 0 | 0 | 0 |
| **Trichomes** | 0 | 207 | 0 | 0 | 0 | 0 | 0 | 0 | 0 |
| **Vascular_Parenchyma** | 9504 | 2516 | 1012 | 551 | 406 | 319 | 0 | 2948 | 2145 |
| **Xylem_Parenchyma** | 2181 | 970 | 173 | 345 | 213 | 584 | 101 | 1186 | 1097 |

Five major cell types were consistently detected across multiple organs: epidermis, marked by cuticle-associated genes such as non-specific lipid transfer proteins, *ECERIFERUM 3-like*, and GDSL esterases; xylem parenchyma, defined by *WAT1*-related proteins, peroxidases, and aquaporins; phloem parenchyma, enriched for sugar transporters and nodulin MtN21 family proteins; companion cells, identified by *PHLOEM PROTEIN 2-LIKE* genes; and a vascular parenchyma population characterized by developmental regulators including homeobox proteins, early nodulin-like genes, and cyclic nucleotide-gated channel proteins.

Meristematic tissue was detected in root tips, dormant buds, and stem cambium (**Fig. 2**), consistent with the known distribution of proliferative zones in perennial plants. In buds, this population further resolved into dormant and proliferating meristem states, reflecting the seasonal transition between dormancy and active growth resumption characteristic of grapevine axillary buds. For the remaining organs (flower, berry, and leaves) no meristematic cluster was identified, consistent with these tissues having been sampled at differentiated developmental stages.

Each organ also contained specialized cell types reflecting its anatomical and developmental context, ranging from 7 cell types in leaves and berries to 16 in flowers. In leaves, mesophyll cells were defined by photosynthetic genes such as *Lhcb1*, and guard cells by the potassium channel *KAT1*. In berries, mesocarp cells were characterized by sucrose-phosphate synthases and bidirectional sugar transporters, while the exocarp was enriched for chalcone synthases. A proliferative green exocarp subpopulation showed co-expression of *cyclin D3.1* and anthocyanin biosynthesis genes, including *flavonoid 3′-hydroxylase* and *glutathione S-transferase F10-like*. Flowers contained diverse anatomical structures and cell types defined by canonical markers, including *POLLENLESS 3-LIKE 2* in microsporangia, *STIG1-like* in stigma, *AGL11* and *AGL6* in ovary, and *SEPALLATA-1* in calyx. It is possible that a greater number of nuclei from each anatomical structure would be required to resolve the individual cell types within anther, stigma, filament, calyx, calyptra and style with greater confidence. Resolving the constituent cell types within these reproductive structures at finer resolution will be an interesting direction for future work. Dormant bud meristematic populations were enriched for histone genes and chromatin modifiers, whereas cortex cells expressed TIFY 5A-like and auxin-responsive regulators. Stems contained vascular cambium (cyclins), protophloem (*SIEVE ELEMENT OCCLUSION B*), protoxylem (*XCP2-like*), and trichomes enriched for terpene synthases such as myrcene synthase and geranyl linalool synthase. Root developmental zones were resolved from meristem (cyclins) through endodermis (*CASP-like protein 1D1*), with trichoblasts distinguished by nutrient transporters and hormone-related genes.

Shared transcriptional features were also observed across anatomically related tissues. For example, cortex cells in dormant buds and stems displayed similar expression profiles, including dormancy-associated regulators and cell wall remodeling enzymes, suggesting a shared regulatory module in these structurally related organs (**Supplementary Text S1**). These observations indicate that some cell types maintain consistent transcriptional identities across organs, whereas others are more strongly shaped by organ context.

To assess whether the cell-type identity programs captured in our atlas reflect deep, conserved regulatory logic rather than grapevine- or perennial-specific features, we tested whether the main marker genes distinguishing companion cells, guard cells, mesophyll, epidermis, and other cell types in our data show the analogous expression pattern in an independent, single-cell-resolved Arabidopsis atlas (Lee et al., 2025), after ortholog assignment via AEGIS (**Table S8**). Two markers replicated consistently: the grapevine companion-cell marker ortholog *g001600*- *AT4G19840* (PHLOEM PROTEIN 2-LIKE family) was highest in the *Arabidopsis* stele, the vascular-bundle-containing tissue where phloem companion cells reside, independently in four organs/developmental stages (21-day rosette, imbibed seed, 6- and 12-day seedlings). The grapevine guard-cell marker ortholog *g219670*-*AT4G24480* (protein kinase superfamily) showed an even more consistent pattern, ranking highest in Arabidopsis guard cells in five of seven organs tested, with the two exceptions (tapetum in flower, cambium in stem) representing biologically plausible alternative deployments in specialized secretory or meristematic contexts rather than random noise (**Fig. S3**). Mesophyll marker orthologs (light-harvesting-complex genes) showed developmentally appropriate specificity: they matched Arabidopsis mesophyll cells precisely in organs with differentiated photosynthetic tissue (mature rosette leaf, seedling) but not in undifferentiated stages (imbibed seed, young rosette) where no such tissue yet exists. In contrast, pollen and xylem-parenchyma marker orthologs did not reproduce their expected cell-type association in Arabidopsis, indicating that not all cell-identity programs captured in our atlas are deeply conserved across the ∼120 million years separating grapevine and Arabidopsis (Fawcett et al., 2009).

### Organ-specific gene regulatory networks

To link TF expression to regulatory programs, we inferred gene regulatory networks by combining FIMO motif scan with MINI-EX2 regulon activity analysis (Staut et al., 2026). We identified a total of 1,971 regulons, each comprising a transcription factor and its predicted target genes (**Table S9**), of which 831 in leaves, 635 in flowers, 273 in berries, 226 in stems and substantially fewer in dormant buds (4) and roots (2).

To systematically compare regulon activity across the atlas, we quantified regulon activity in all cells using AUCell (Aibar et al., 2017), averaged activity scores by organ, cell type, and replicate, and partitioned variance using a linear mixed model. Organ identity explained the largest share of regulon activity variance (65 ± 24 %), followed by residual noise (18 ± 14 %) and cell type (17 ± 19 %) (**Fig. 3A; Fig. S4**). Applying an adjusted p-value (p-adj) threshold of 1 x 10⁻⁸, we identified organ-enriched regulons in each organ: 517 in flowers, 276 in young leaves, 222 in senescent leaves, 446 in green berries, 55 in harvest berries, 188 in postharvest berries, 41 in stems, 7 in buds, and 142 in roots (**Fig. 3B**).

**Figure 3.**
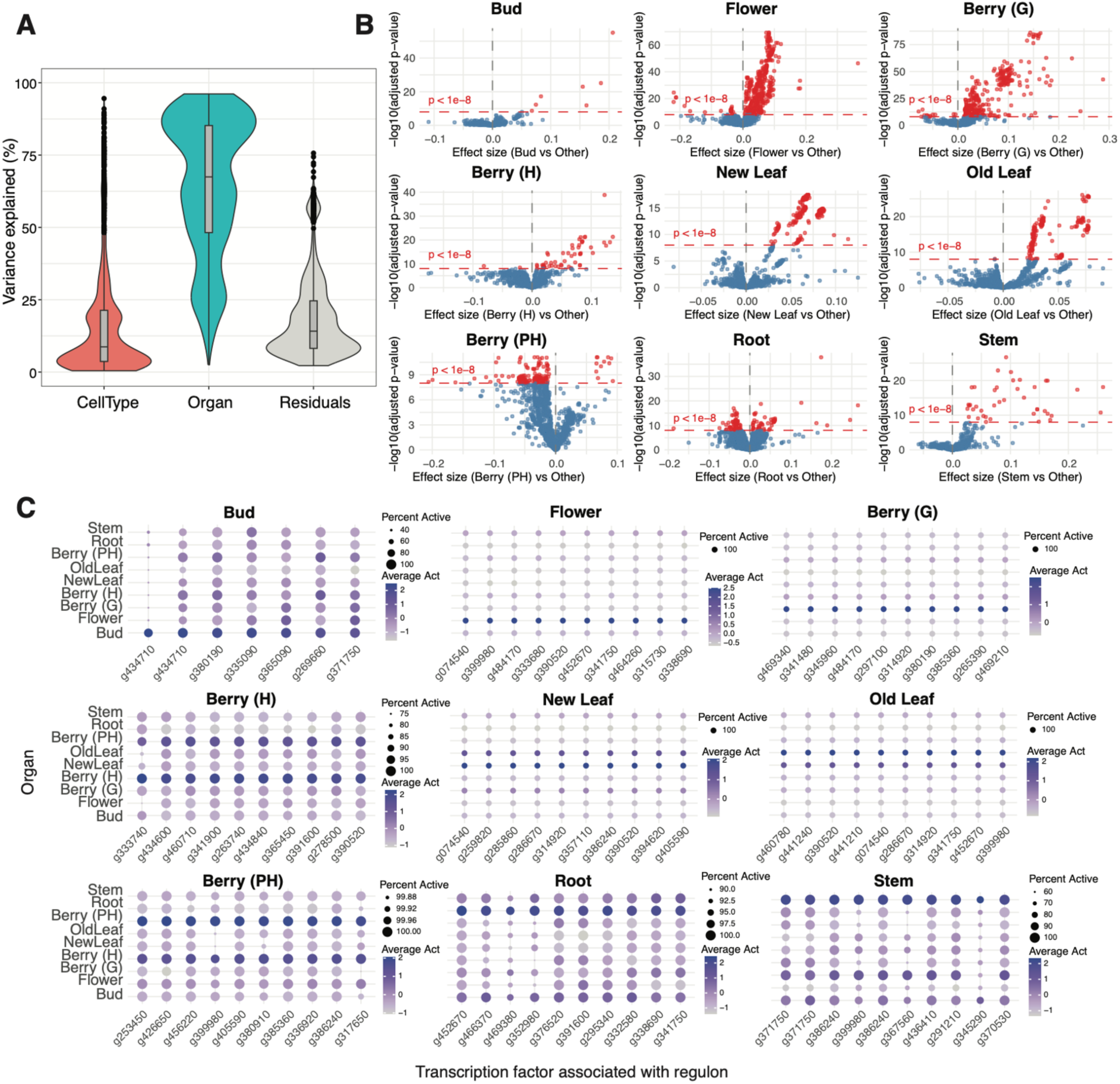
Organ-specific gene regulatory network activity across the grapevine atlas. (A) Partitioning of variance in regulon activity across the atlas, showing the relative contributions of organ, cell type, and residual variation based on a linear mixed model, with organ and cell type as fixed effects and biological replicate as a random effect. (B) Volcano plots of regulon activity differences for each organ compared to all other organs, showing effect size versus -log10(p-adj). The dashed line indicates the significance threshold (p-adj < 1 × 10⁻⁸), used to identify organ-enriched regulons. (C) Dot plots of the top 10 organ-enriched regulons for each organ, showing average regulon activity (color scale) and percentage of nuclei with active regulons (dot size) across organs.

Several organ-enriched regulons had clear biological relevance. The regulon driven by a no apical meristem-like (*NAM*) family protein (*g456220*) was enriched in postharvest berries and included target genes involved in sugar metabolism (*g324860*) and transport (*g451000*), and ripening (**Table S10**), consistent with the known NAM role in fruit maturation (Liu et al., 2023). A regulon driven by *YABBY2* (*g341480*) was enriched in green berries and targeted five aquaporins, over a total of 60 genes, pointing to a role in cell expansion and water transport (**Fig. 3C**; **Table S9-10**), consistent with its role in early berry maturation (Shen et al., 2024). A regulon driven by the transcription factor *GATA15* (*g295340*) was enriched in roots and included chromatin and cell wall remodeling, consistent with its role in lateral root formation (Kiryushkin et al., 2019).

While most regulons showed organ-specific activity, a subset was specifically active in companion cells across organs, pointing to a regulatory architecture that is maintained independently of organ context, as described in the following sections.

### Integrated atlas reveals organ-dependent transcriptional organization

Across the integrated dataset, Louvain-based clusters 5 and 24 contained more than 80% of all annotated companion cells (8,561 of 10,778; **Fig. 4A-B**). Clusters 5 and 24 were enriched for canonical companion cell markers including *PHLOEM PROTEIN 2*, alongside genes supporting their metabolic functions, including raffinose and galactinol synthases (sugar metabolism and transport), *UDP-glucose/UDP-galactose 4-epimerase* (cell wall biosynthesis), and *beta-amylase* (starch degradation and sugar mobilization). Companion cells were further enriched for genes involved in redox homeostasis and detoxification (isoflavone reductase-like, thioredoxin H2, copper chaperone), calcium signaling (*CML7, CML17, CML19, CML29*), and metal homeostasis (heavy metal transport/detoxification proteins and cation/H⁺ antiporters) (**Table S11**), along with several uncharacterized genes that represent candidate regulators of companion cell function.

**Figure 4.**
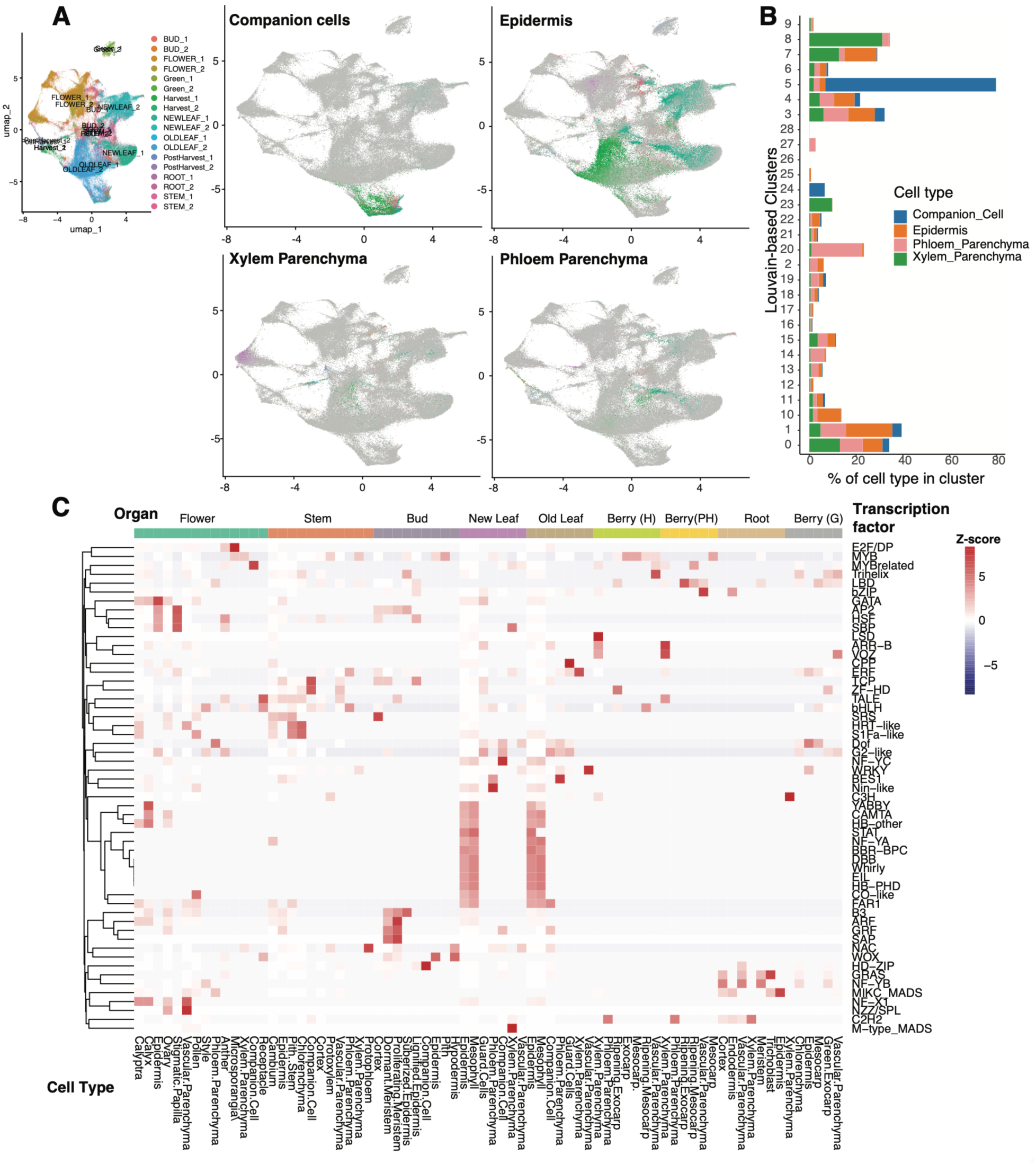
Integrated multi-organ single-nucleus transcriptomic atlas of grapevine. (A) UMAP highlighting the distribution of selected shared cell types (companion cells, phloem parenchyma, xylem parenchyma, and epidermis) across organs. (B) Distribution of major shared cell types across Louvain clusters, showing the proportion of each cell type within clusters. (C) Heatmap of aggregated transcription factor family expression (Z-scores) across cell types and organs, revealing cell type-and organ-associated regulatory patterns.

By contrast, other cell types shared across organs, including epidermis, xylem parenchyma, and phloem parenchyma, did not form unified clusters across the atlas but instead segregated according to organ context (**Fig. 4A-B**). These results indicate that, for most cell types, organ identity dominates transcriptional variation, whereas in companion cells, cell identity is maintained largely independent of organ context.

### Cell type and organ-specific transcription factor expression

To further characterize the transcriptional landscape of the atlas, we next examined transcription factor (TF) expression across organs and cell types. Using PlantTFDB (Jin et al., 2016), we catalogued homologous TF genes between PN40024 and Pixie. In total, 1,196 TFs were identified, of which 882 were expressed in the atlas. The highest number of highly expressed TFs was observed in mesophyll and epidermal cells of leaves, followed by the flower ovary and stem cambium (**Table S12**). Several TFs showed highly restricted expression, including a bZIP TF (*g255780*) in the root endodermis, a nodulation-signaling pathway 2 protein-like TF (*g293350*) in root trichoblasts, a growth-regulating factor 10-like TF (*g264030*) in the ovary, and three lateral organ boundaries (LOB) domain TFs in pollen. TF families also displayed cell type and organ-specific patterns (**Fig. 4C**); for example, E2F/DP family TFs, core regulators of the cell cycle (Miao et al., 2024), were highly expressed in flower microsporangia (**Fig. 4C**).

### Whole-plant cell type specific gene regulatory networks

From the linear mixed model, we identified 711 cell type-specific regulons (p-adj < 1 × 10^-8^; **Fig. 5A**). The most distinct cell type was microsporangium, with 184 enriched regulons, followed by xylem parenchyma (162), pollen (143), companion cells (57), anther (41), epidermis (31), and other cell types with fewer enriched regulons (listed in **Table S9-10**).

**Figure 5.**
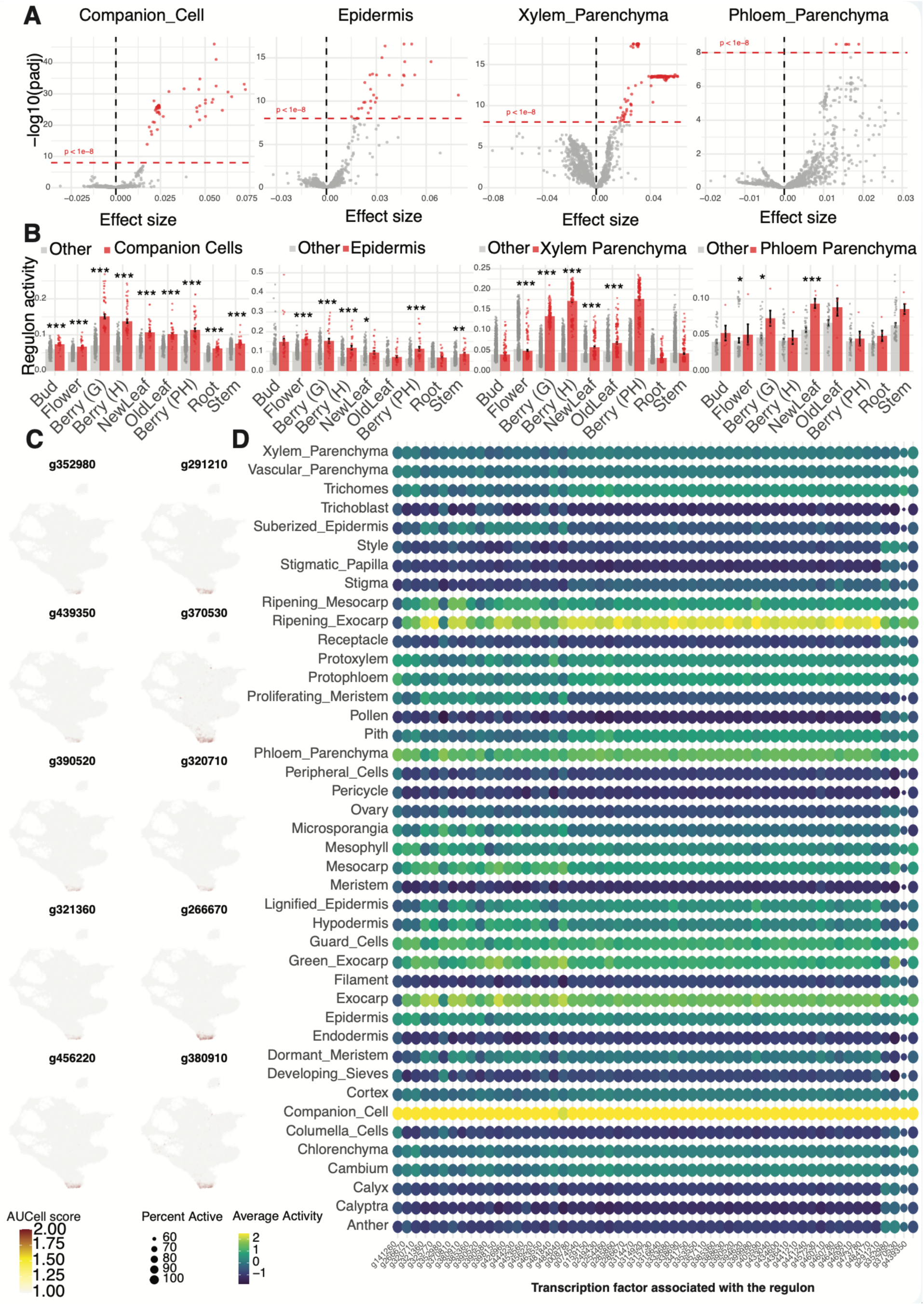
Cell type-specific gene regulatory network activity across the grapevine atlas. (A) Volcano plots showing differential regulon activity for major cell types shared across organs (companion cells, epidermis, xylem parenchyma, and phloem parenchyma), comparing each cell type to all others. Effect size is plotted against -log10(p-adj); the dashed line indicates the significance threshold (p-adj < 1 × 10⁻⁸). (B) Regulon activity distributions across organs for each focal cell type, compared to all other cell types. Bars represent mean regulon activity ± SE; points correspond to individual nuclei. Asterisks indicate significant differences relative to other cell types (* p-adj < 0.05; ** < 0.01; *** < 0.001). (C) UMAP feature plots showing activity of representative companion cell-active regulons across the atlas. The gene name of the transcription factor associated with the regulon is noted on top of each UMAP. (D) Dot plot of regulon activity across cell types. Color indicates average regulon activity and dot size the proportion of nuclei with active regulons. Only regulons significantly enriched in companion cells are shown.

Among cell types shared across organs, including epidermis, xylem parenchyma, phloem parenchyma, and companion cells, only companion cells showed consistently enriched regulons when compared to all other cell types across all organs (**Table S10**; **Fig. 5B**). This pattern is consistent with the organ-independent clustering of companion cells in the integrated atlas (**Fig. 4A**), and together these results support a regulatory architecture in companion cells for which the same regulons are independently and repeatedly identified as cell-type-specific across all sampled organs. Nonetheless, the absolute magnitude of regulon activity in companion cells varied significantly among organs (Fig. S5, Table S14), consistent with differences in overall transcriptomic composition and sequencing depth across organs, which affect the absolute scale of an AUCell score independently of cell-type identity and are therefore not informative about companion cell specificity per se.

Notable regulons consistently more active in companion cells across organs were driven by the TFs *g320710* (DNA polymerase epsilon subunit 3), *g321360* and *g380910* (bHLH family), *g390520* and *g370530* (ERFs), *g291210* (B-box zinc finger), *g352980* and *g439350* (MYB-related), and *g456220* (no apical meristem-like) (**Fig. 5C**). Target genes spanned transport, metabolism, signaling, and redox regulation (**Table S13**), including sucrose transporters (*g226840*, *g472820*), amino acid permeases (*g338460*), aquaporins, ion transporters, vesicle trafficking components, proton pumps, and pyrophosphate-energized vacuolar proton pumps (**Fig. 5D**; **Table S12**) consistent with the energetic demands of phloem loading (Otero and Helariutta, 2016). Regulons were further enriched for redox homeostasis genes including peroxidases, thioredoxins, nucleoredoxins, and cytochrome-related proteins, alongside *WRKY* TFs, bHLH-like proteins, RNA-binding proteins, and hormone- and calcium-related targets including auxin-and gibberellin-responsive genes and *CML* proteins (**Table S13**).

Together, these companion cell regulons define a coherent and organ-independent transcriptional architecture centered on phloem loading, redox balance, and signaling, distinguishing companion cells from other shared cell types that undergo extensive organ-specific transcriptional reprogramming.

### Cell type specific developmental trajectories

Having defined cell types across nine organs, we next asked whether the atlas could resolve developmental dynamics within individual cell types. To test this, we reconstructed developmental trajectories for pollen germination, berry mesocarp and exocarp maturation, and leaf mesophyll and epidermis senescence by redefining clusters within each cell type and applying pseudotime analysis using Monocle 3 (Trapnell et al., 2014). Gene markers, refined cluster annotation and nuclei count per developmental stage are summarized in **Table S15A-C**.

For pollen, clusters were assigned to immature (33), maturing (1), mature (10, 11, 37, 40), and germinating (23) states. Pseudotime increased progressively from immature to germinating cells (**Fig. 6A-B**). Genes associated with this trajectory included a *GRAS* transcription factor, aquaporins, *SNAP25*-like proteins, and pectate lyase (**Fig. 6C**), reflecting a transition from early differentiation to hydration, cell wall remodeling, and vesicle trafficking during germination.

**Figure 6.**
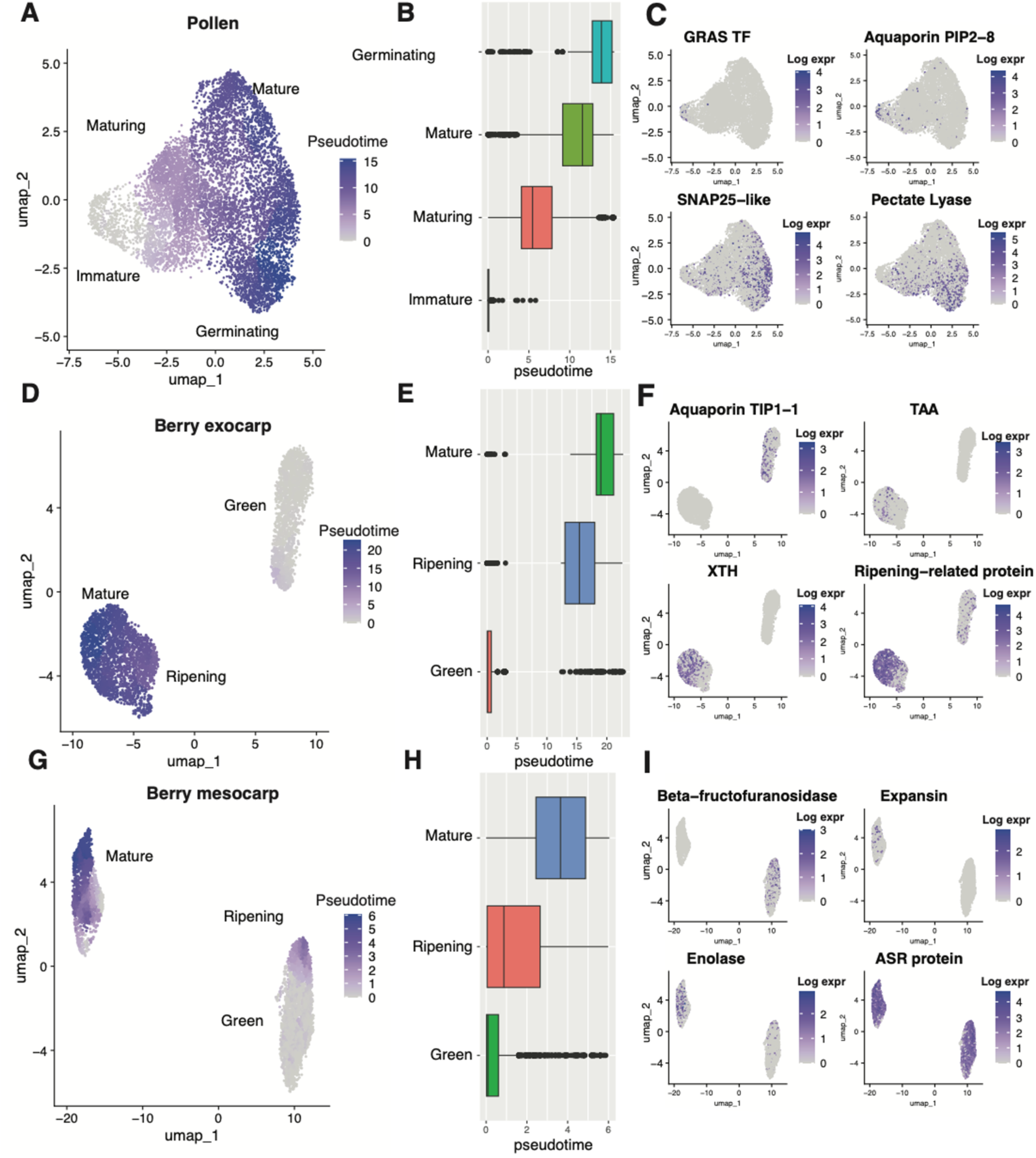
Cell type-specific developmental trajectories across grapevine organs. UMAP embeddings colored by pseudotime illustrate developmental progression inferred using Monocle 3 for pollen (A), berry exocarp (D), and berry mesocarp (G). Corresponding boxplots summarize pseudotime distributions across annotated developmental stages for each cell type (B, E, H), confirming the progression from early to late states. Feature plots show log-normalized expression of genes significantly associated with pseudotime (q-value < 0.05) across each trajectory (C, F, I), highlighting coordinated transcriptional changes during pollen germination and berry tissue maturation.

For berry exocarp, clusters were classified as green (1, 13), ripening (8, 12, 14), and mature (4). Pseudotime increased from green to mature stages (**Fig. 6D-E**). Associated genes included *TIP1-1*, tryptophan aminotransferase (*TAA*), *XTH*, and ripening-related proteins (**Fig. 6F**), reflecting a shift from cell expansion to cell wall remodeling and ripening-associated processes (**Table S16**).

For berry mesocarp, clusters were classified as green (3, 7), ripening (2), and mature (5) stages. Pseudotime increased from green to mature mesocarp (**Fig. 6G-H**). Associated genes included *β-fructofuranosidase*, expansins, enolases, and abscisic stress-related (*ASR*) proteins (**Fig. 6I**), reflecting a transition from primary carbon metabolism and sugar accumulation to cell wall modification and ripening-associated processes (**Table S16**).

For leaf epidermis, clusters were grouped into young (3, 22), mature (4, 12, 18, 24), and senescent (6, 10, 11, 15, 16) stages. Pseudotime increased toward senescence (**Fig. S6A-C**). Associated genes included endoglucanases, *FERONIA*, *GAPC*, and Octicosapeptide/Phox/Bem1p family proteins (**Fig. S6C**), reflecting a shift from active metabolism to cell wall remodeling and senescence-associated responses (**Table S16**).

For leaf mesophyll, clusters were defined as young (2), mature (1, 17), and senescent (0, 19, 20). Pseudotime increased toward senescence (**Fig. S6B-D**). Associated genes included photosystem II components (*psbY*), aquaporin *PIP1-2*, UDP-glucosyltransferases, and *STAY-GREEN* proteins (**Fig. S6F**), reflecting a transition from photosynthetic activity to stress response and senescence-associated metabolic reprogramming (**Table S16**).

Together, these trajectories confirm that the atlas resolves transcriptional dynamics at sub-organ resolution across diverse developmental processes, establishing a foundation for examining how cell identity is maintained or remodeled across organ contexts.

### Transcriptional plasticity across shared cell types

To compare organ-specific transcriptional variation across shared cell types, we subset companion cells, epidermis, xylem parenchyma, and phloem parenchyma from the integrated atlas, renormalized the data, and performed pseudobulk differential expression analysis across organs. Genes with p-adj < 0.05 were considered differentially expressed, and Z-scores were used to assess the magnitude and direction of expression differences. The resulting datasets comprised 10,778 companion cell nuclei expressing 26,661 genes (**Fig. 7A**), 48,535 epidermal nuclei expressing 36,344 genes (**Fig. 7C**), 6,850 xylem parenchyma nuclei expressing 26,692 genes (**Fig. 7E**), and 6,339 phloem parenchyma nuclei expressing 25,074 genes (**Fig. 7G**).

**Figure 7.**
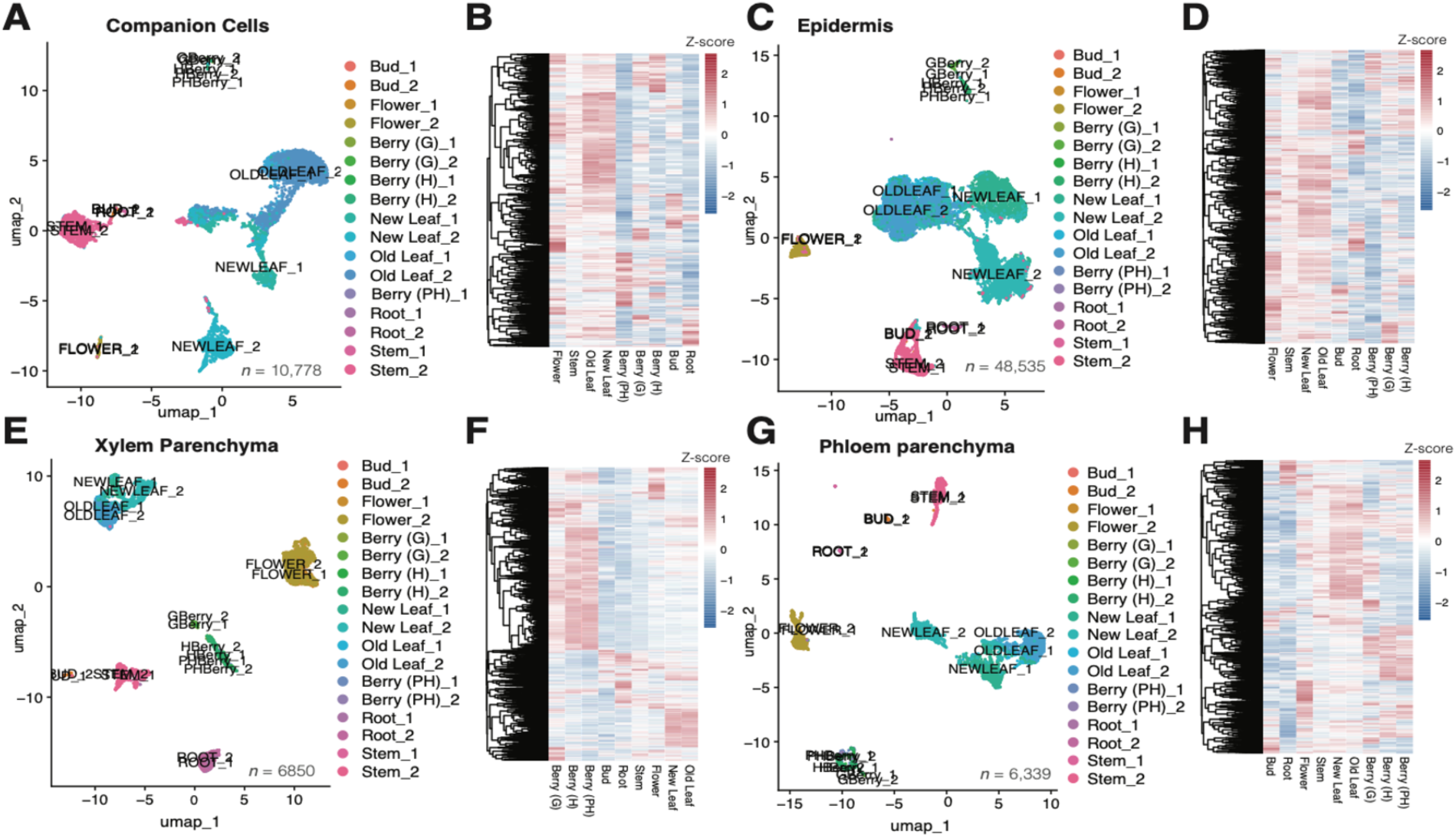
Organ-associated transcriptional variation in shared cell types. (A, C, E, G) UMAP embeddings of nuclei from companion cells (A), epidermis (C), xylem parenchyma (E), and phloem parenchyma (G), colored by organ and biological replicate. (B, D, F, H) Heatmaps showing z-scored expression of differentially expressed genes (likelihood ratio test, p-adj < 0.05) across organs for each corresponding cell type. Rows represent genes and columns represent organs; hierarchical clustering highlights organ-associated expression patterns.

Companion cells showed the fewest DEGs across organs (3,223; 12% of expressed genes) (**Fig. 7B**), followed by phloem parenchyma (4,231; 17%) (**Fig. 7H**), xylem parenchyma (11,255; 42%) (**Fig. 7F**), and epidermis (14,824; 41%) (**Fig. 7D**). This gradient of transcriptional plasticity is consistent with the regulon and clustering analyses: companion cells maintain a stable core transcriptional program with organ-associated shifts reflecting local physiological roles, including stress and dormancy responses in buds, sugar transport and ripening in berries, and water and carbon import in roots (**Table S17**). In contrast, epidermis and xylem parenchyma undergo extensive organ-specific reprogramming. Epidermal cells transition from protective and stress-related programs in buds to reproductive specialization in flowers, water transport and cuticle formation in developing fruits, pigmentation and ripening-associated metabolism linked to seed dispersal, and senescence-associated degradation in aging leaves (**Table S18**). Xylem parenchyma shifts between stress tolerance, defense, transport, and metabolic programs depending on organ and developmental stage (**Table S19**). Phloem parenchyma occupies an intermediate position, with organ-specific adaptations that partially parallel companion cell programs, particularly in sugar transport, stress responses, and cell wall remodeling (**Table S20**). Detailed pseudobulk results for all shared cell types are provided in **Supplementary Text S2**.

Together, these results define a continuum of transcriptional behavior across shared cell types, from highly constrained in companion cells to highly plastic in epidermis and xylem parenchyma, with phloem parenchyma at an intermediate position. This continuum is consistent with the variance partitioning of regulon activity, in which organ identity accounted for 65 ± 24% of total variance compared to 17 ± 19% for cell type (**Fig. 3A**), confirming that organ context is the dominant driver of transcriptional variation across most shared cell types, with companion cells as a notable exception.

Finally, the τ specificity index was applied to organ-level pseudobulk profiles (averaged across all cells per organ, excluding organellar transcripts). A total of 210 organ-specific genes (τ > 0.85) across all nine organs was identified, and their annotations recapitulated known organ biology closely (**Table S21**). Flower (57 genes) was dominated by reproductive hallmarks (myrcene synthase, pollen-specific extensin, and pollen-tube RALF signaling peptide), as previously reported (Fasoli et al., 2012). Young and senescent leaf formed a clean developmental contrast: active photosynthetic machinery (photosystem I subunits, plastocyanin) versus the canonical senescence regulator STAY-GREEN. Root was marked by aquaporin PIP1-1 and the dehydration-responsive gene RD22, matching its water-uptake/stress-sensing role. The berry samples tracked ripening biology, with harvest berry dominated by thaumatin-like proteins and a β-1,3-glucanase, among the most well-documented ripening markers in grape (Fasoli et al., 2012). Most notably, bud yielded a coherent cold-hardiness and dormancy signature (dehydrin, Dehydrin Rab15, a low-temperature-induced protein, a prion-like stress protein), in line with previous bulk transcriptomic approaches (Fasoli et al., 2012). Stem yielded the fewest organ-exclusive genes (5).

## Discussion

This study presents a multi-organ single-nucleus transcriptomic atlas of a woody perennial crop, integrating more than 220,000 nuclei across nine organs and 46 annotated cell types and anatomical structures. Comparable multi-organ atlases have been generated in Arabidopsis, soybean, and rice (Guo et al., 2025; Lee et al., 2025; Yan et al., 2025; Zhang et al., 2025), but not in woody perennials. Grapevine captures features not represented in annual models, including extensive secondary growth, woody vasculature, and perennial-specific cell types such as ray parenchyma, vessel-associated cells, and dormant bud meristematic populations. The atlas also spans developmental transitions that are rarely present in annual systems, including dormancy, budbreak, and a berry developmental series spanning green to postharvest stages. Together, these features extend multi-organ single-cell frameworks to perennial plant biology.

Cross-referencing our results with the previously published Arabidopsis single-nucleus transcriptomics atlas (Lee et al., 2025), we identified companion-cell and guard-cell marker genes representing an evolutionarily ancient, broadly conserved regulatory core (**Fig S3**), consistent with our finding that companion cells maintain organ-independent transcriptional programs within grapevine itself, while other cell-type-specific expression signatures are more evolutionarily and developmentally labile. Future studies should be conducted to confirm this hypothesis.

The cultivar Pixie was selected because it provides a particularly useful experimental system for grapevine functional genomics, owing to its compact architecture and ability to flower and fruit year-round under greenhouse conditions. Beyond providing a comprehensive reference of cellular diversity across grapevine organs, this atlas provides a foundation for investigating the cellular basis of the Pixie dwarf phenotype. Comparisons with closely related cultivars, such as Pinot Meunier, could help identify cell types, developmental programs, and molecular pathways associated with the compact growth habit and accelerated reproductive development characteristic of Pixie.

Several cell-type marker genes identified in our atlas were consistent with markers reported in previous grapevine single-cell and single-nucleus studies, providing additional support for the robustness of our cell-type annotations. For example, *ECERIFERUM* was associated with epidermal cells, *PHOTOSYSTEM II SUBUNIT O* with mesophyll cells, and *Cyclin A1* and members of the Histone superfamily with proliferating meristematic cells (Bonarota et al., 2025b; Li et al., 2026; Liang et al., 2026; Yao et al., 2026). The recurrence of these markers across independent grapevine single-cell datasets suggests that these transcriptional signatures capture conserved cellular programs and provides confidence in the biological interpretation of the cell populations resolved in our atlas.

Most plant single-cell studies profile individual organs, limiting the ability to separate cell-intrinsic transcriptional programs from those shaped by organ context; profiling nine organs within a single framework overcomes this limitation. We observed a continuum in which some cell types maintain similar transcriptional profiles across organs, whereas others are extensively reshaped by organ context. Variance partitioning of regulon activity quantifies this pattern, with organ identity explaining 65 ± 24% of the variance compared to 17 ± 19% for cell type. Thus, for most shared cell types, organ context is the dominant driver of transcriptional variation. Gene regulatory networks are thus extensively reconfigured by organ context in most cell types, rather than operating as fixed modules. This quantitative framework for classifying cell types by their degree of context dependence extends concepts emerging from multi-organ plant atlases (Guo et al., 2025; Lee et al., 2025) to a perennial species.

Within this continuum, companion cells stand out as the most transcriptionally stable shared cell type, supported by three independent observations: organ-independent clustering in the integrated atlas (**Fig. 4A**), the lowest number of differentially expressed genes among shared cell types (**Table S17**), and a set of conserved regulons active across organs (**Fig. 5**). Companion cells, together with sieve elements, play a central role in plant productivity by partitioning assimilates between source and sink organs (Khadilkar et al., 2016), yet their metabolic functions and regulatory control remain incompletely understood (Otero and Helariutta, 2016). Their primary role is to sustain enucleated sieve elements, supporting phloem loading and unloading of sugars and the transport of signaling molecules, including hormones, proteins, and RNA (Matilla, 2023). This functional specialization imposes consistent constraints across organs. In contrast to cell types such as the epidermis, which adopt distinct roles depending on organ context, companion cells appear to operate under similar physiological demands throughout the plant (Oparka and Turgeon, 1999; Turgeon and Wolf, 2009; Van Bel, 2003). The conserved regulons identified here, enriched for transport, redox balance, and signaling functions, are consistent with this role. In particular, phloem loading is driven by proton gradients at the companion cell plasma membrane (Taiz and Zeiger, 2010), generated by proton-pumping ATPases and pyrophosphatases (H⁺-PPi) (Hunt et al., 2023). We identified two H⁺-PPi genes (*g137540* and *g383290*) that are consistently expressed in companion cells across organs and belong to a regulon controlled by the B-box zinc finger TF *g291210*, previously implicated in stress responses (Aydemir et al., 2020). Whether this stability is conserved across plant species or reflects grapevine-specific phloem organization remains open questions that comparative single-cell studies in Arabidopsis and other species could directly address (Guo et al., 2025;Lee et al., 2025).

In contrast to companion cells, epidermis and xylem parenchyma showed the highest DEG counts among shared cell types (**Fig. 7B, F**) and extensive organ-specific transcriptional reprogramming. This divergence reflects the inherently context-dependent nature of their core functions, with dominant transcriptional programs shifting across organs rather than being uniformly maintained. In dormant buds, expression is enriched for stress-associated pathways and lipid metabolism linked to surface barrier reinforcement. In flowers, it is dominated by regulators of reproductive development and volatile biosynthesis. In mature berries, the program is centered on pigmentation, defense, and ripening-associated metabolism, while in senescing leaves it shifts toward degradation pathways and nutrient remobilization (Javelle et al., 2011; Yeats and Rose, 2013). Xylem parenchyma shows a similar pattern of reprogramming, but along a different functional axis. Across organs, its transcriptional profile alternates between stress tolerance, transport, and secondary metabolism, with no single program consistently dominant across all organs. This is consistent with its role at the interface of transport and metabolism, where functional demands vary with organ identity and developmental stage (Morris et al., 2016; Secchi et al., 2017). Phloem parenchyma occupies an intermediate position, with organ-specific shifts that partially mirror companion cell programs, suggesting that proximity to the sieve element–companion cell complex imposes constraints on the extent of reprogramming.

Pseudotime analysis resolved developmental dynamics within individual cell types across three agronomically important processes: pollen maturation, berry ripening, and leaf senescence. These transitions underlie fruit set, photosynthetic capacity, and yield, yet their cell type-specific genetic regulation has remained poorly characterized due to the lack of cellular resolution in bulk approaches (Wei et al., 2010). Pollen trajectories captured a progression from early differentiation to hydration, vesicle trafficking, and cell wall remodeling during germination. Grapevine pollen is binucleate at anthesis (Mićić et al., 2018) and possesses a transcriptional profile geared toward rapid resource mobilization and energy production (Fasoli et al., 2012); the upregulation of 17 zinc finger proteins in germinating pollen further supports the involvement of this gene family in grapevine pollen development (Arrey-Salas et al., 2021). Berry exocarp and mesocarp showed distinct maturation programs: exocarp trajectories were associated with auxin signaling, cell wall remodeling, and ripening-associated processes (Klee and Giovannoni, 2011), while mesocarp maturation was driven by sugar metabolism, ABA signaling, and cell expansion consistent with the metabolic shift of ripening (Lijavetzky et al., 2012). Leaf senescence trajectories distinguished epidermal and mesophyll programs: mesophyll senescence was marked by the decline of photosystem II components and a shift toward UDP-glucosyltransferase activity (Guo et al., 2021; Vangelisti et al., 2020), while epidermal senescence was characterized by FERONIA and cell wall remodeling genes, reflecting a distinct tissue-specific degradation program. These trajectories reinforce the broader pattern observed across the atlas, in which developmental progression is accompanied by coordinated, cell type-specific reprogramming, further highlighting the extent of transcriptional plasticity outside of conserved cell types such as companion cells.

While these analyses provide cell type resolution of developmental programs, several limitations should be considered. Single-nucleus sequencing captures nuclear transcripts and may underrepresent cytoplasmic or rapidly turned-over mRNAs, potentially limiting detection of genes with low nuclear localization (Bakken et al., 2018; Grindberg et al., 2013). In addition, cell type annotation across organs relies on marker conservation and may be affected by context-dependent shifts in marker gene expression. The regulon predictions are based on motif inference and co-expression and require chromatin-level validation through ATAC-seq or DAP-seq to confirm TF binding *in vivo* (Bajic et al., 2017; Farmer et al., 2021; Feng et al., 2022; Zhang et al., 2026). The atlas represents a single cultivar under greenhouse conditions, and transcriptional programs may differ in field-grown vines or across *Vitis* diversity. Future work integrating epigenomic data, cross-cultivar sampling, and functional validation of candidate regulators will be needed to fully exploit the regulatory framework established here.

Despite these limitations, this atlas establishes a whole-plant, cell type resolved reference for grapevine and provides a framework for understanding how gene regulatory networks are organized across organs in a perennial crop. The central finding, that transcriptional plasticity varies systematically across shared cell types, with companion cells as a stable regulatory anchor and epidermis and xylem parenchyma as highly plastic extremes, reframes cell identity as a balance between conserved regulatory cores and context-dependent reprogramming across organs. The resource, the regulatory inferences, and the conceptual framework together provide a foundation for dissecting development, stress responses, and vascular function in grapevine and other woody perennials.

## Author contribution

M.-S.B. and D.C. conceptualized and designed the project. D.C. supervised the project and acquired funding. M.-S.B. performed all experimental work including sampling and nuclei extractions. R.F.-B. contributed to nuclei isolation. M.-S.B. performed the data analyses and visualization. N.C. contributed to bioinformatic analyses. M.-S.B. and D.C. wrote the manuscript.

## Data availability

Sequencing data are accessible through NCBI under the BioProject PRJNA1457451. The seurat object including the raw and normalized data, cell type annotation, and regulon activity is available in Zenodo (https://zenodo.org/records/22739600). An interactive website was created to explore this dataset (https://www.grapegenomics.com/pages/VvPixie/Pixie_snRNAseq/atlas_snrnaseq.php).

## Code availability

The analysis pipeline used in this study is available at https://github.com/mariasoleb/Pixie_snRNAseq_atlas

## Acknowledgments

We thank Hong Qiu (UC Davis) for the 10x Genomics library preparation and the UC Davis DNA Technologies and Expression Analysis Core Facility for sequencing. This work was partially funded by the USDA NIFA Award 2022-51181-38240, the USDA-ARS cooperative agreement #58-2030-5-076, the E.J. Gallo Winery, and the Ray Rossi Endowment in Viticulture and Enology.

## References

Aibar, S., González-Blas, C.B., Moerman, T., Huynh-Thu, V.A., Imrichova, H., Hulselmans, G., Rambow, F., Marine, J.-C., Geurts, P., Aerts, J., 2017. SCENIC: single-cell regulatory network inference and clustering. Nature methods 14, 1083–1086.

Arrey-Salas, O., Caris-Maldonado, J.C., Hernández-Rojas, B., Gonzalez, E., 2021. Comprehensive genome-wide exploration of C2H2 zinc finger family in grapevine (Vitis vinifera L.): Insights into the roles in the pollen development regulation. Genes 12, 302.

Arroyo-García, R., Ruiz-García, L., Bolling, L., Ocete, R., López, M., Arnold, C., Ergul, A., Söylemezo Lu, G., Uzun, H., Cabello, F., 2006. Multiple origins of cultivated grapevine (Vitis vinifera L. ssp. sativa) based on chloroplast DNA polymorphisms. Molecular ecology 15, 3707–3714.

Badia-i-Mompel, P., Wessels, L., Müller-Dott, S., Trimbour, R., Ramirez Flores, R.O., Argelaguet, R., Saez-Rodriguez, J., 2023. Gene regulatory network inference in the era of single-cell multi-omics. Nat Rev Genet 24, 739–754. 10.1038/s41576-023-00618-5

Bailey, T.L., Johnson, J., Grant, C.E., Noble, W.S., 2015. The MEME suite. Nucleic acids research 43, W39–W49.

Bajic, M., Maher, K.A., Deal, R.B., 2017. Identification of open chromatin regions in plant genomes using ATAC-Seq, in: Plant Chromatin Dynamics: Methods and Protocols. Springer, pp. 183–201.

Bakken, T.E., Hodge, R.D., Miller, J.A., Yao, Z., Nguyen, T.N., Aevermann, B., Barkan, E., Bertagnolli, D., Casper, T., Dee, N., 2018. Single-nucleus and single-cell transcriptomes compared in matched cortical cell types. PloS one 13, e0209648.

Baumgart, L.A., Greenblum, S.I., Morales-Cruz, A., Wang, P., Zhang, Y., Yang, L., Chen, C., Dilworth, D.J., Garretson, A.C., Grosjean, N., 2025. Recruitment, rewiring and deep conservation in flowering plant gene regulation. Nature Plants 1–14.

Blanc-Mathieu, R., Dumas, R., Turchi, L., Lucas, J., Parcy, F., 2024. Plant-TFClass: a structural classification for plant transcription factors. Trends in plant science 29, 40–51.

Bonarota, M.-S., Garcia, J.F., Massonnet, M., Zaccheo, M., Figueroa-Balderas, R., Cochetel, N., Cantu, D., 2025. Dual single-nucleus gene expression atlas of grapevine and Erysiphe necator during early powdery mildew infection. Molecular Plant-Microbe Interactions MPMI-08.

Boss, P.K., Thomas, M.R., 2002. Association of dwarfism and floral induction with a grape ‘green revolution’ mutation. Nature 416, 847–850. 10.1038/416847a

Braun, D.M., 2022. Phloem loading and unloading of sucrose: what a long, strange trip from source to sink. Annual Review of Plant Biology 73, 553–584.

Çakır Aydemir, B., Yüksel Özmen, C., Kibar, U., Mutaf, F., Büyük, P.B., Bakır, M., Ergül, A., 2020. Salt stress induces endoplasmic reticulum stress-responsive genes in a grapevine rootstock. PLoS One 15, e0236424.

Cantó-Pastor, A., Kajala, K., Shaar-Moshe, L., Manzano, C., Timilsena, P., De Bellis, D., Gray, S., Holbein, J., Yang, H., Mohammad, S., 2024. A suberized exodermis is required for tomato drought tolerance. Nature plants 10, 118–130.

Cantu, D., Massonnet, M., Cochetel, N., 2024. The wild side of grape genomics. Trends in Genetics 40, 601–612.

Cao, X., Ma, T., Fan, R., Yuan, G.-C., 2024. Systematic analysis identifies a connection between spatial and genomic variations of chromatin states. Cell systems 15, 1092–1102.

Castro-Mondragon, J.A., Riudavets-Puig, R., Rauluseviciute, I., Berhanu Lemma, R., Turchi, L., Blanc-Mathieu, R., Lucas, J., Boddie, P., Khan, A., Manosalva Pérez, N., 2022. JASPAR 2022: the 9th release of the open-access database of transcription factor binding profiles. Nucleic acids research 50, D165–D173.

Chen, Y., Tong, S., Jiang, Y., Ai, F., Feng, Y., Zhang, J., Gong, J., Qin, J., Zhang, Y., Zhu, Y., Liu, J., Ma, T., 2021. Transcriptional landscape of highly lignified poplar stems at single-cell resolution. Genome Biol 22, 319. 10.1186/s13059-021-02537-2

Chen, Y.-L., Hsieh, J.-W.A., Kuo, S.-C., Kao, C.-T., Tung, C.-C., Yu, J.-H., Chang, T.-H., Ku, C., Xie, J., Zhang, D., Li, Q., Lin, Y.-C.J., 2024. Merit of integrating in situ transcriptomics and anatomical information for cell annotation and lineage construction in single-cell analyses of Populus. Genome Biol 25, 85. 10.1186/s13059-024-03227-5

Cochetel, N., Minio, A., Guarracino, A., Garcia, J.F., Figueroa-Balderas, R., Massonnet, M., Kasuga, T., Londo, J.P., Garrison, E., Gaut, B.S., 2023. A super-pangenome of the North American wild grape species. Genome Biology 24, 290.

Conde, D., Triozzi, P.M., Pereira, W.J., Schmidt, H.W., Balmant, K.M., Knaack, S.A., Redondo-López, A., Roy, S., Dervinis, C., Kirst, M., 2022. Single-nuclei transcriptome analysis of the shoot apex vascular system differentiation in Populus. Development 149, dev200632. 10.1242/dev.200632

Cousins, P., 2012. Small but Mighty: “Pixie” Grapevine Speeds Up the Pace of Grape Genetics Research and Breeding. Cornell’s Viticulture and Enology 2:1−4.

Denyer, T., Ma, X., Klesen, S., Scacchi, E., Nieselt, K., Timmermans, M.C.P., 2019. Spatiotemporal Developmental Trajectories in the Arabidopsis Root Revealed Using High-Throughput Single-Cell RNA Sequencing. Developmental Cell 48, 840–852.e5. 10.1016/j.devcel.2019.02.022

Dorrity, M.W., Cuperus, J.T., Carlisle, J.A., Fields, S., Queitsch, C., 2018. Preferences in a trait decision determined by transcription factor variants. Proceedings of the National Academy of Sciences 115, E7997–E8006.

Farmer, A., Thibivilliers, S., Ryu, K.H., Schiefelbein, J., Libault, M., 2021. Single-nucleus RNA and ATAC sequencing reveals the impact of chromatin accessibility on gene expression in Arabidopsis roots at the single-cell level. Molecular plant 14, 372–383.

Fasoli, M., Dal Santo, S., Zenoni, S., Tornielli, G.B., Farina, L., Zamboni, A., Porceddu, A., Venturini, L., Bicego, M., Murino, V., 2012. The grapevine expression atlas reveals a deep transcriptome shift driving the entire plant into a maturation program. The Plant Cell 24, 3489–3505.

Fawcett, J.A., Maere, S., Van De Peer, Y., 2009. Plants with double genomes might have had a better chance to survive the Cretaceous–Tertiary extinction event. Proceedings of the National Academy of Sciences 106, 5737–5742.

Feng, D., Liang, Z., Wang, Y., Yao, J., Yuan, Z., Hu, G., Qu, R., Xie, S., Li, D., Yang, L., 2022. Chromatin accessibility illuminates single-cell regulatory dynamics of rice root tips. BMC biology 20, 274.

Goremykin, V.V., Salamini, F., Velasco, R., Viola, R., 2009. Mitochondrial DNA of Vitis vinifera and the issue of rampant horizontal gene transfer. Molecular biology and evolution 26, 99–110.

Grant, C.E., Bailey, T.L., Noble, W.S., 2011. FIMO: scanning for occurrences of a given motif. Bioinformatics 27, 1017–1018.

Grindberg, R.V., Yee-Greenbaum, J.L., McConnell, M.J., Novotny, M., O’Shaughnessy, A.L., Lambert, G.M., Araúzo-Bravo, M.J., Lee, J., Fishman, M., Robbins, G.E., 2013. RNA-sequencing from single nuclei. Proceedings of the National Academy of Sciences 110, 19802–19807.

Grones, C., Eekhout, T., Shi, D., Neumann, M., Berg, L.S., Ke, Y., Shahan, R., Cox, K.L., Gomez-Cano, F., Nelissen, H., Lohmann, J.U., Giacomello, S., Martin, O.C., Cole, B., Wang, J.-W., Kaufmann, K., Raissig, M.T., Palfalvi, G., Greb, T., Libault, M., De Rybel, B., 2024. Best practices for the execution, analysis, and data storage of plant single-cell/nucleus transcriptomics. Plant Cell 36, 812–828. 10.1093/plcell/koae003

Guo, X., Wang, Yichuan, Zhao, C., Tan, C., Yan, W., Xiang, S., Zhang, D., Zhang, H., Zhang, M., Yang, L., Yan, M., Xie, P., Wang, Yi, Li, L., Fang, D., Guang, X., Shao, W., Wang, F., Wang, H., Sahu, S.K., Liu, M., Wei, T., Peng, Y., Qiu, Y., Peng, T., Zhang, Y., Ni, X., Xu, Z., Lu, H., Li, Z., Yang, H., Wang, E., Lisby, M., Liu, H., Guo, H., Xu, X., 2025. An Arabidopsis single-nucleus atlas decodes leaf senescence and nutrient allocation. Cell 188, 2856–2871.e16. 10.1016/j.cell.2025.03.024

Guo, Y., Ren, G., Zhang, K., Li, Z., Miao, Y., Guo, H., 2021. Leaf senescence: progression, regulation, and application. Molecular Horticulture 1, 5.

Harrison Day, B.L., Brodersen, C.R., Brodribb, T.J., 2024. Weak link or strong foundation? Vulnerability of fine root networks and stems to xylem embolism. New Phytologist 244, 1288–1302.

He, D., Patro, R., 2023. simpleaf: a simple, flexible, and scalable framework for single-cell data processing using alevin-fry. Bioinformatics 39, btad614.

He, D., Zakeri, M., Sarkar, H., Soneson, C., Srivastava, A., Patro, R., 2022. Alevin-fry unlocks rapid, accurate and memory-frugal quantification of single-cell RNA-seq data. Nature Methods 19, 316–322.

Healy, J., McInnes, L., 2024. Uniform manifold approximation and projection. Nature Reviews Methods Primers 4, 82.

Honma, T., Goto, K., 2001. Complexes of MADS-box proteins are sufficient to convert leaves into floral organs. Nature 409, 525–529. 10.1038/35054083

Hunt, H., Brueggen, N., Galle, A., Vanderauwera, S., Frohberg, C., Fernie, A.R., Sonnewald, U., Sweetlove, L.J., 2023. Analysis of companion cell and phloem metabolism using a transcriptome-guided model of Arabidopsis metabolism. Plant physiology 192, 1359– 1377.

Illouz-Eliaz, N., Yu, J., Swift, J., Lande, K., Jow, B., Partida-Garcia, L., Tuang, Z.K., Lee, T.A., Yaaran, A., Gomez-Castanon, R., 2025. Drought recovery in plants triggers a cell-state-specific immune activation. Nature Communications 16, 8095.

Javelle, M., Vernoud, V., Rogowsky, P.M., Ingram, G.C., 2011. Epidermis: the formation and functions of a fundamental plant tissue. New Phytologist 189, 17–39.

Jin, J., Tian, F., Yang, D.-C., Meng, Y.-Q., Kong, L., Luo, J., Gao, G., 2016. PlantTFDB 4.0: toward a central hub for transcription factors and regulatory interactions in plants. Nucleic acids research gkw982.

Khadilkar, A.S., Yadav, U.P., Salazar, C., Shulaev, V., Paez-Valencia, J., Pizzio, G.A., Gaxiola, R.A., Ayre, B.G., 2016. Constitutive and companion cell-specific overexpression of AVP1, encoding a proton-pumping pyrophosphatase, enhances biomass accumulation, phloem loading, and long-distance transport. Plant physiology 170, 401–414.

Kim, J.-Y., Symeonidi, E., Pang, T.Y., Denyer, T., Weidauer, D., Bezrutczyk, M., Miras, M., Zöllner, N., Hartwig, T., Wudick, M.M., 2021. Distinct identities of leaf phloem cells revealed by single cell transcriptomics. The Plant Cell 33, 511–530.

Kiryushkin, A.S., Ilina, E.L., Puchkova, V.A., Guseva, E.D., Pawlowski, K., Demchenko, K.N., 2019. Lateral root initiation in the parental root meristem of cucurbits: Old players in a new position. Frontiers in Plant Science 10, 365.

Klee, H.J., Giovannoni, J.J., 2011. Genetics and control of tomato fruit ripening and quality attributes. Annual review of genetics 45, 41–59.

Kubo, M., Udagawa, M., Nishikubo, N., Horiguchi, G., Yamaguchi, M., Ito, J., Mimura, T., Fukuda, H., Demura, T., 2005. Transcription switches for protoxylem and metaxylem vessel formation. Genes Dev. 19, 1855–1860. 10.1101/gad.1331305

Lee, H.-C., Lin, T.-Y., 2005. Isolation of plant nuclei suitable for flow cytometry from recalcitrant tissue by use of a filtration column. Plant Molecular Biology Reporter 23, 53–58.

Lee, Travis A., Illouz-Eliaz, N., Nobori, T., Xu, J., Jow, B., Nery, J.R., Ecker, J.R., 2025. A single-cell, spatial transcriptomic atlas of the Arabidopsis life cycle. Nat. Plants 11, 1960–1975. 10.1038/s41477-025-02072-z

Levine, M., Davidson, E.H., 2005. Gene regulatory networks for development. Proceedings of the National Academy of Sciences 102, 4936–4942.

Li, H., Dai, X., Huang, X., Xu, M., Wang, Q., Yan, X., Sederoff, R.R., Li, Q., 2021. Single-cell RNA sequencing reveals a high-resolution cell atlas of xylem in Populus. Journal of Integrative Plant Biology 63, 1906–1921. 10.1111/jipb.13159

Li, Y., Bian, Y., Ni, Y., Xie, Z., 2026. Single-nucleus transcriptomic atlas reveals autophagy and hormone signaling in early grapevine (Vitis vinifera L.) tendril versus inflorescence determination. The Plant Journal 125, e70804.

Liang, C., Li, X., He, J., Jin, Y., Wang, R., Song, Y., Shi, X., Du, G., Wang, L., 2026. Single-nucleus transcriptome analysis reveals spatial and cell-type-specific responses to sunburn in grape berries. Horticulture Research uhag310.

Lijavetzky, D., Carbonell-Bejerano, P., Grimplet, J., Bravo, G., Flores, P., Fenoll, J., Hellín, P., Oliveros, J.C., Martinez-Zapater, J.M., 2012. Berry flesh and skin ripening features in Vitis vinifera as assessed by transcriptional profiling. PloS one 7, e39547.

Liu, J., Qiao, Y., Li, C., Hou, B., 2023. The NAC transcription factors play core roles in flowering and ripening fundamental to fruit yield and quality. Frontiers in Plant Science 14, 1095967.

Liu, Zhongjie, Wang, N., Su, Y., Long, Q., Peng, Y., Shangguan, L., Zhang, F., Cao, S., Wang, X., Ge, M., Xue, H., Ma, Z., Liu, W., Xu, X., Li, C., Cao, X., Ahmad, B., Su, X., Liu, Y., Huang, G., Du, M., Liu, Zhenya, Gan, Y., Sun, L., Fan, X., Zhang, C., Zhong, H., Leng, X., Ren, Y., Dong, T., Pei, D., Wu, X., Jin, Z., Wang, Y., Liu, C., Chen, J., Gaut, B., Huang, S., Fang, J., Xiao, H., Zhou, Y., 2024. Grapevine pangenome facilitates trait genetics and genomic breeding. Nat Genet 56, 2804–2814. 10.1038/s41588-024-01967-5

Love, M.I., Huber, W., Anders, S., 2014. Moderated estimation of fold change and dispersion for RNA-seq data with DESeq2. Genome biology 15, 1–21.

Lun, A., Griffiths, J., McCarthy, D., He, D., Patro, R., 2018. DropletUtils: utilities for handling single-cell droplet data. R package version 0.99 14.

Marand, A.P., Chen, Z., Gallavotti, A., Schmitz, R.J., 2021. A *cis*-regulatory atlas in maize at single-cell resolution. Cell 184, 3041–3055.e21. 10.1016/j.cell.2021.04.014

Matilla, A.J., 2023. The interplay between enucleated sieve elements and companion cells. Plants 12, 3033.

Miao, Y., You, H., Liu, H., Zhao, Y., Zhao, J., Li, Y., Shen, Y., Tang, D., Liu, B., Zhang, K., 2024. RETINOBLASTOMA RELATED 1 switches mitosis to meiosis in rice. Plant Communications 5.

Mićić, N., Đurić, G., Jovanović Cvetković, T., Cvetković, M., 2018. Pollen functional ability in two indigenous grapevine cultivars in Bosnia and Herzegovina. Eur J Hortic Sci 83, 35– 41.

Morris, H., Brodersen, C., Schwarze, F.W., Jansen, S., 2016. The parenchyma of secondary xylem and its critical role in tree defense against fungal decay in relation to the CODIT model. Frontiers in Plant Science 7, 1665.

Navarro-Payá, D., Santiago, A., Velt, A., Moretto, M., Rustenholz, C., Matus, J.T., 2026. AEGIS: an annotation extraction and genomic integration resource. Bioinformatics 42, btag363.

Oparka, K.J., Turgeon, R., 1999. Sieve elements and companion cells—traffic control centers of the phloem. The Plant Cell 11, 739–750.

Otero, S., Helariutta, Y., 2016. Companion cells: a diamond in the rough. Journal of experimental botany erw392.

Paineau, M., Liang, C., Cochetel, N., Shivani, Figueroa-Balderas, R., Thilmony, R., Rumbaugh, A., Cantu, D., 2025. Integrative genome and transcriptome analysis identifies smoke-responsive glycosyltransferases in grapevine berries. Journal of Experimental Botany 76, 6032–6050.

Satija, R., Farrell, J.A., Gennert, D., Schier, A.F., Regev, A., 2015. Spatial reconstruction of single-cell gene expression data. Nature biotechnology 33, 495–502.

Secchi, F., Pagliarani, C., Zwieniecki, M.A., 2017. The functional role of xylem parenchyma cells and aquaporins during recovery from severe water stress. Plant, cell & environment 40, 858–871.

Shan, H., Cheng, J., Zhang, R., Yao, X., Kong, H., 2019. Developmental mechanisms involved in the diversification of flowers. Nature plants 5, 917–923.

Shen, H., Luo, B., Ding, Y., Xiao, H., Chen, G., Yang, Z., Hu, Z., Wu, T., 2024. The YABBY transcription factor, SLYABBY2A, positively regulates fruit septum development and ripening in tomatoes. International Journal of Molecular Sciences 25, 5206.

Shulse, C.N., Cole, B.J., Ciobanu, D., Lin, J., Yoshinaga, Y., Gouran, M., Turco, G.M., Zhu, Y., O’Malley, R.C., Brady, S.M., Dickel, D.E., 2019. High-Throughput Single-Cell Transcriptome Profiling of Plant Cell Types. Cell Reports 27, 2241–2247.e4. 10.1016/j.celrep.2019.04.054

Staut, J., Pérez, N.M., Depuydt, T., Vandepoele, K., Lukicheva, S., 2026. MINI-EX version 2: cell-type-specific gene regulatory network inference using an integrative single-cell transcriptomics approach, in: Plant Transcription Factors: Methods and Protocols. Springer, pp. 159–191.

Taiz, L., Zeiger, E., 2010. Plant physiology 5 th (Ed.). Sunderland. Sinauer Assoc. Inc., Publishers, Sunderland Masschusetts.

Tang, H., Krishnakumar, V., Zeng, X., Xu, Z., Taranto, A., Lomas, J.S., Zhang, Y., Huang, Y., Wang, Y., Yim, W.C., 2024. JCVI: A versatile toolkit for comparative genomics analysis. Imeta 3, e211.

Tenorio Berrío, R., Verhelst, E., Eekhout, T., Grones, C., De Veylder, L., De Rybel, B., Dubois, M., 2025. Dual and spatially resolved drought responses in the Arabidopsis leaf mesophyll revealed by single-cell transcriptomics. New Phytologist 246, 840–858.

Theißen, G., Melzer, R., Rümpler, F., 2016. MADS-domain transcription factors and the floral quartet model of flower development: linking plant development and evolution. Development 143, 3259–3271. 10.1242/dev.134080

Trapnell, C., Cacchiarelli, D., Grimsby, J., Pokharel, P., Li, S., Morse, M., Lennon, N.J., Livak, K.J., Mikkelsen, T.S., Rinn, J.L., 2014. The dynamics and regulators of cell fate decisions are revealed by pseudotemporal ordering of single cells. Nature biotechnology 32, 381–386.

Turgeon, R., Wolf, S., 2009. Phloem transport: cellular pathways and molecular trafficking. Annual review of plant biology 60, 207–221.

Van Bel, A., 2003. The phloem, a miracle of ingenuity. Plant, Cell & Environment 26, 125–149.

Vangelisti, A., Guidi, L., Cavallini, A., Natali, L., Lo Piccolo, E., Landi, M., Lorenzini, G., Malorgio, F., Massai, R., Nali, C., 2020. Red versus green leaves: transcriptomic comparison of foliar senescence between two Prunus cerasifera genotypes. Scientific Reports 10, 1959.

Wang, Y., Huan, Q., Li, K., Qian, W., 2021. Single-cell transcriptome atlas of the leaf and root of rice seedlings. Journal of Genetics and Genomics 48, 881–898. 10.1016/j.jgg.2021.06.001

Wang, Y., Tang, H., Wang, X., Sun, Y., Joseph, P.V., Paterson, A.H., 2024. Detection of colinear blocks and synteny and evolutionary analyses based on utilization of MCScanX. Nature Protocols 19, 2206–2229.

Wei, L.Q., Xu, W.Y., Deng, Z.Y., Su, Z., Xue, Y., Wang, T., 2010. Genome-scale analysis and comparison of gene expression profiles in developing and germinated pollen in Oryza sativa. Bmc Genomics 11, 338.

Yamaguchi, M., Kubo, M., Fukuda, H., Demura, T., 2008. VASCULAR-RELATED NAC-DOMAIN7 is involved in the differentiation of all types of xylem vessels in Arabidopsis roots and shoots. The Plant Journal 55, 652–664. 10.1111/j.1365-313X.2008.03533.x

Yan, H., Mendieta, J.P., Zhang, X., Luo, Z., Marand, A.P., Liang, Y., Minow, M.A., Zhong, Y., Jin, Y., Jang, H., 2025. A single-cell rice atlas integrates multi-species data to reveal cis-regulatory evolution. Nature Plants 11, 2050–2071.

Yanai, I., Benjamin, H., Shmoish, M., Chalifa-Caspi, V., Shklar, M., Ophir, R., Bar-Even, A., Horn-Saban, S., Safran, M., Domany, E., 2005. Genome-wide midrange transcription profiles reveal expression level relationships in human tissue specification. Bioinformatics 21, 650–659.

Yao, X., Xu, Z., Xi, Y., He, X., Gao, Q., Lu, J., Fu, P., 2026. Unveiling the early defense response dynamics in grapevines against Plasmopara viticola by single-cell transcriptomics. Genome Biology 27, 43.

Yeats, T.H., Rose, J.K., 2013. The formation and function of plant cuticles. Plant physiology 163, 5–20.

Zhang, T.-Q., Chen, Y., Liu, Y., Lin, W.-H., Wang, J.-W., 2021. Single-cell transcriptome atlas and chromatin accessibility landscape reveal differentiation trajectories in the rice root. Nature communications 12, 2053.

Zhang, X., Luo, Z., Marand, A.P., Yan, H., Jang, H., Bang, S., Mendieta, J.P., Minow, M.A.A., Schmitz, R.J., 2025. A spatially resolved multi-omic single-cell atlas of soybean development. Cell 188, 550–567.e19. 10.1016/j.cell.2024.10.050

Zhang, Y., Liu, H., Xu, M., Xie, M., Deng, S., Ma, X., Zhao, L., He, F., Li, M., Long, R., 2026. Single-nucleus RNA-seq and ATAC-seq analyses provide molecular insights into the cadmium stress response in alfalfa roots. Horticulture Research uhag117.

Zhu, A., Srivastava, A., Ibrahim, J.G., Patro, R., Love, M.I., 2019. Nonparametric expression analysis using inferential replicate counts. Nucleic Acids Research 47, e105–e105.

